# Fibrin is a critical regulator of neutrophil effector function at mucosal barrier sites

**DOI:** 10.1101/2021.01.15.426743

**Authors:** Lakmali M Silva, Andrew D Doyle, Collin L Tran, Teresa Greenwell-Wild, Nicolas Dutzan, Andrew G Lum, Cary S Agler, Megan Sibree, Priyam Jani, Daniel Martin, Vardit Kram, Francis J Castellino, Matthew J Flick, Kimon Divaris, Thomas H Bugge, Niki M Moutsopoulos

## Abstract

Tissue-specific cues are critical for homeostasis at mucosal barriers. Here, we document that the clotting factor fibrin is a critical regulator of neutrophil function at mucosal barriers. We demonstrate that fibrin engages neutrophils through the α_M_β_2_ integrin receptor and activates effector functions, including the production of reactive oxygen species and NET formation. These immune-protective neutrophil functions become tissue damaging in the context of impaired plasmin-mediated fibrinolysis. Indeed, the accumulation of fibrin due to Mendelian genetic defects in plasmin leads to severe oral mucosal immunopathology in mice and humans. Concordantly, genetic polymorphisms in the human *PLG* gene, encoding plasminogen, are associated with common forms of the oral mucosal disease periodontitis. Our work uncovers fibrin as a critical regulator of neutrophil effector function within the mucosal tissue microenvironment and suggests fibrin-neutrophil engagement as a pathogenic instigator and therapeutic target in common mucosal disease.

## Introduction

Mucosal sites of the human body are poised to encounter continuous stimulation from the environment and must balance outside stimuli without overexerting inflammation while providing protection from infection and injury [1]. Indeed, mucosal tissue-specific immune responses are key for the regulation of tissue homeostasis but when disrupted will lead to disease [1, 2]. Despite well-established appreciation of this general concept, specific mechanisms of mucosal immunity are not yet completely understood. Furthermore, the relevance of specific pathways in human mucosal immunity to susceptibility of mucosal disease is often difficult to determine. However, Mendelian diseases can be instrumental towards understanding human biology. Specifically, Mendelian diseases can uncover genes and pathways implicated in human immunity and mucosal homeostasis. One Mendelian disease clearly linked to mucosal disease is that of Plasminogen deficiency. Plasminogen (Plg), the zymogen precursor of the serine protease plasmin, is synthesized by the liver and is present in high concentration in the circulation and in interstitial fluids to provide focal proteolysis, after being converted to plasmin by urokinase or tissue plasminogen activator. Homozygous or compound heterozygous mutations in the human Plg gene *(PLG)* typically lead to severe mucosal disease involvement suggesting, a critical role for this gene/pathway in mucosal immunity. Indeed, such patients present with deposition of fibrin at various mucosal sites leading to ocular disease (conjunctivitis), oral mucosal disease (ligneous periodontitis), lung, vaginal and gastrointestinal tract involvement [3–5]. In the oral mucosa, local deposition of fibrin is hypothesized to lead to severe soft tissue and bone destruction around teeth and often loss of the entire dentition in adolescence [6–9]. Mucosal inflammation in areas surrounding the dentition and destruction of underlying bone are also the hallmarks of the common human oral mucosal disease, periodontitis [10, 11].

Herein, we hypothesized that insufficient plasmin-mediated fibrinolysis is a key contributor to mucosal immunopathology in humans and in animal models. To understand the mechanistic links between mucosal fibrin deposition and immunopathology, an array of genetically engineered mouse models complemented by histological and genetic studies in humans were employed. We chose to study the mechanisms involved in Plg deficiency-mediated immunopathology at the site of the oral mucosa, as the oral disease periodontitis is amongst the most common manifestations in humans with Plg deficiency, suggesting that this mucosal site is particularly susceptible to Plg deficiency-related immunopathology. Our work highlights the role of fibrin as a critical immune-regulator of mucosal barrier homeostasis and provides insights into the role of fibrin in neutrophil activation and tissue immunity.

## Results

### Plg deficiency instigates oral mucosal immunopathology in mice

*Plg^-/-^* mice phenocopy Plg-deficient humans and present with mucosal immunopathology including periodontitis, conjunctivitis and inflammation in the G.I. tract [12–15]. The hallmark of periodontitis is mucosal inflammation and destruction of bone around teeth. Indeed, *Plg^-/-^* mice develop spontaneous periodontal bone loss by 12 weeks of age when compared to their wild-type and heterozygous littermates, which is increasingly severe at the age of 24 weeks (Figure 1A-B and Supplementary Figure 1A). Consistent with accelerated periodontal bone loss in *Plg^-/-^* mice, mice double deficient for the two physiological plasminogen activators, tPA and uPA (*Plat^-/-^;Plau^-/-^* mice) also display spontaneous severe periodontal bone loss (Supplementary Figure 1B). To further confirm that spontaneous periodontitis in *Plg^-/-^* mice is associated with the loss of enzymatic activity of plasmin, we used a knock-in mouse model that harbors a mutation in the catalytic serine residue (S743A), rendering plasmin catalytically inactive (*Plg^S743A/S743A^*). At the ages of 12 and 24 weeks, *Plg^S743A/S743A^* mice also show a significant increase in periodontal bone loss compared to their wild-type and heterozygous littermate controls, confirming that plasmin-insufficiency drives periodontal bone loss (Figure 1C). Consistent with mucosal lesions in patients with *PLG* deficiency, oral mucosal tissues of *Plg^-/-^* mice display a disruption of tissue architecture associated with deposition of a matrix protein, presumably fibrin(ogen) (Figure 1D and F). Our findings confirm that Plg deficiency in mice, consistent with the human disease, leads to severe oral mucosal immunopathology, which is associated with fibrin(ogen) deposition.

**Figure 1:**
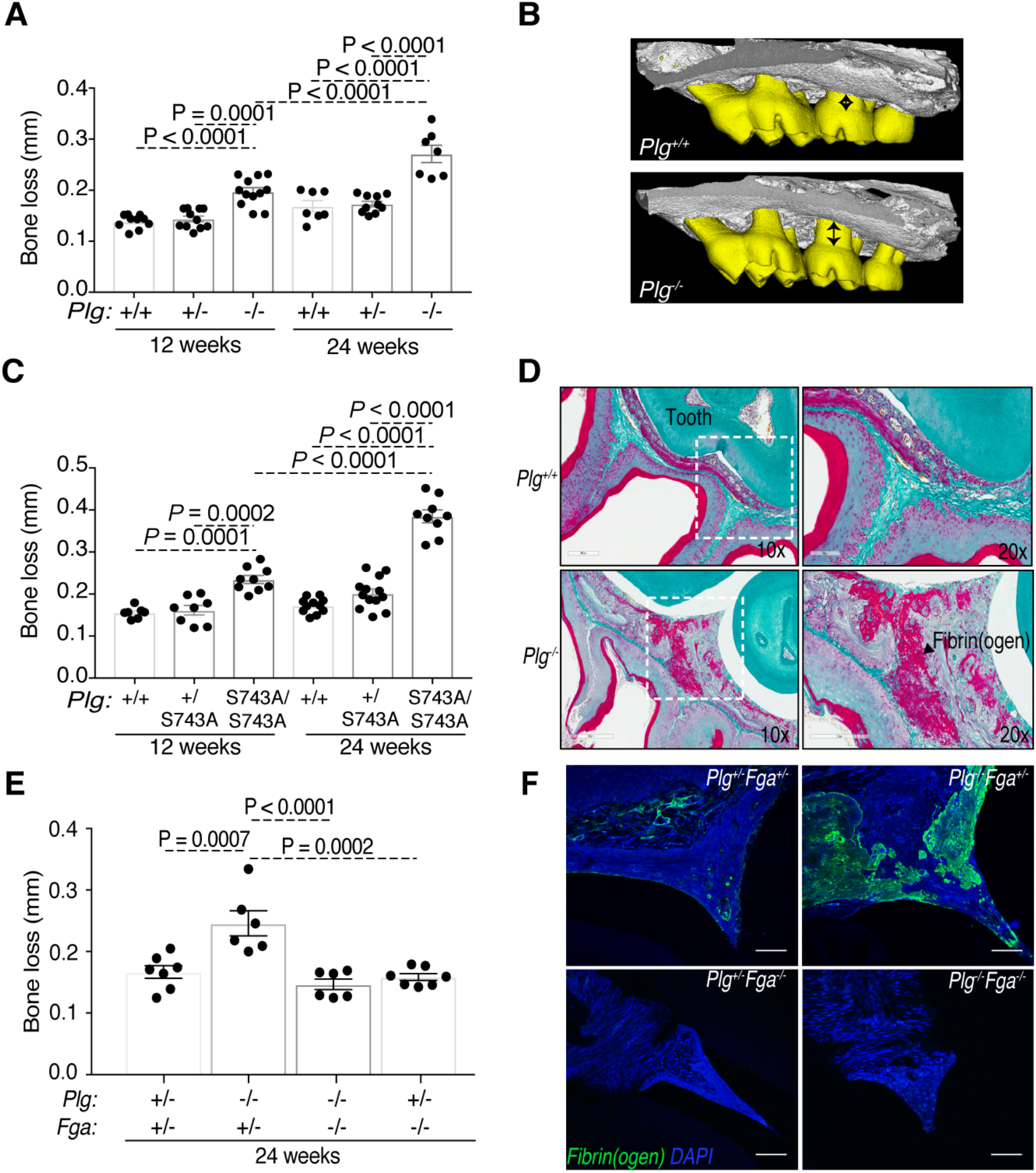
Defective fibrinolysis triggers oral mucosal immunopathology. (A) Plg-deficient mice (*Plg^-/-^)* display spontaneous periodontal bone loss at the age of 12 and 24 weeks compared with *Plg^+/-^* and *Plg^+/+^* littermates. (B) μCT visualization of periodontal bone loss in *Plg^-/-^* and *Plg^+/+^* maxilla (black arrows depict the distance between alveolar bone crest and cemento-enamel junction). (C) Mice with catalytically inactive plasmin (*Plg^S743A/S743A^*) develop significantly increased spontaneous periodontal bone loss. (D) Fraser-Lendrum staining of *Plg^+/+^* and *Plg^-/-^* oral mucosal tissue sections (Green: Collagen; Magenta: keratin, fibrin; and orange: erythrocytes; black arrow depicts fibrin deposition) (E) Bone loss measurements in *Plg^+/-^ Fga^+/-^, Plg^-/-^ Fga^+/-^, Plg^+/-^ Fga^-/-^* and *Plg^-/-^ Fga^-/-^* mouse maxillae. (F) Immunofluorescence (IF) staining for fibrin(ogen) (green) and DAPI (blue) in *Plg^+/+^, Plg^-/-^* and *Fga^-/-^* and *Plg^-/-^ Fga^-/-^* oral tissue sections (20x). Scale bar = 50 μm.

### Elimination of fibrinogen rescues oral mucosal immunopathology in Plg-deficient mice

Based on these findings, we next sought to determine whether fibrin accumulation, secondary to impaired plasmin-mediated fibrinolysis, is the trigger for mucosal immunopathology in Plg deficiency. For this purpose, we generated Plg and fibrinogen double-deficient (*Plg^-/-^;Fga^-/-^*) mice. We examined whether fibrinogen deficiency would prevent the oral mucosal phenotype in *Plg^-/-^* mice. As expected, double-deficient (*Plg^-/-^;Fga^-/-^*) mice did not demonstrate fibrin deposition in mucosal tissues, contrary to their *Plg^-/-^* littermates (Figure 1F). Importantly, fibrinogen deficiency completely prevented the periodontal bone loss imposed by Plg deficiency (Figure 1E). These data demonstrate that oral mucosal immunopathology in Plg deficiency is indeed driven by fibrin(ogen) accumulation.

### Oral mucosal lesions in Plg-deficient mice are neutrophil-dominated

We next aimed to dissect the mechanisms of fibrin-mediated mucosal immunopathology. To this end, we took an unbiased approach and evaluated global transcriptomic changes by RNA sequencing of oral mucosal (gingival) tissues of *Plg^-/-^* mice compared to wild-type littermates. Principal component analysis of RNA-seq results revealed a distinct transcriptome in mucosal tissues of *Plg^-/-^* mice (Figure 2A). Gene Ontology (GO) analysis of the most significantly upregulated pathways in Plg deficiency revealed neutrophil migration as the top differentially expressed pathway, followed by inflammation, extracellular matrix disassembly, cytokine signaling, leukocyte migration and response to bacterial molecules (Figure 2B). Evaluation of the most significantly differentially expressed genes between *Plg^-/-^* and wild-type mice (Figure 2C), revealed upregulation of genes associated with neutrophil recruitment and granulopoiesis, including chemokine (C-X-C motif) ligand (*Cxcl*) *1-3* and *5,* colony stimulating factor 3 (*Csf3*) and Csf3 receptor (*Csf3r*). Genes involved in neutrophil/monocyte activation were also upregulated, including pro-platelet basic protein (*Ppbp*), ADAM metallopeptidase domain 8 (*Adam8),* triggering receptor expressed on myeloid cells 1 (*Trem1*), C-type lectin domain family 4, member E (*Clec4e*) and CD300 molecule-like family member B (*Cd300lb*) (Figure 2D). Finally, several matrix metalloproteases (*Mmp8-10* and *Mmp12*), interleukins (*Il1α* and *Il1β)* and prostaglandin-endoperoxide synthase 2 (*Ptgs2*) that are involved in extracellular matrix remodeling and inflammation are also upregulated in *Plg^-/-^* mice. These data reveal an upregulation of inflammatory pathways that is dominated by a neutrophil signature.

**Figure 2:**
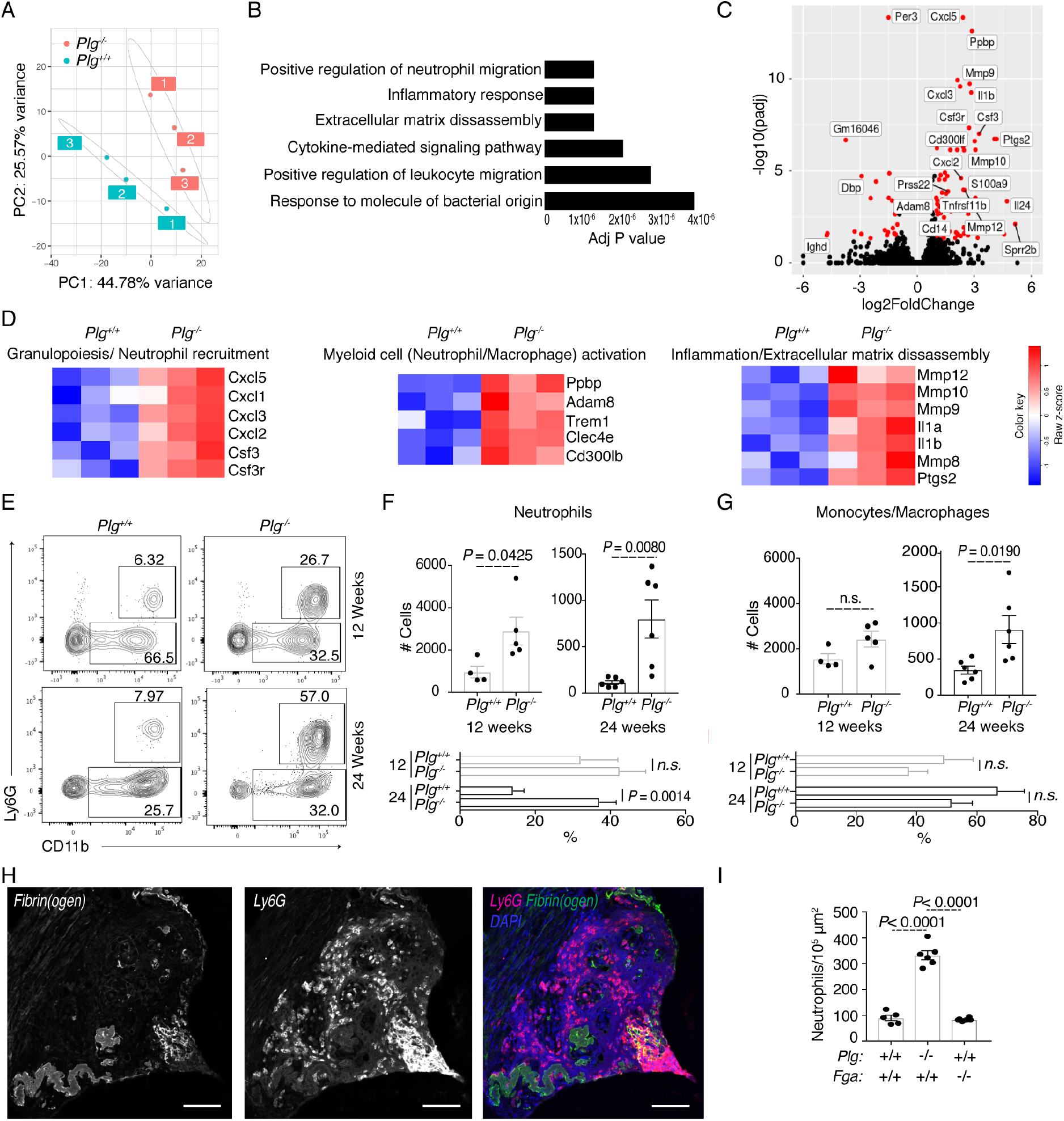
Lesions of mucosal immunopathology in Plg-deficiency are dominated by neutrophils. (A) PCA plot showing the distribution of differentially expressed genes in oral mucosa of 12-week-old *Plg^+/+^* and *Plg^-/-^* mice. (B) Gene Ontology (GO) – biological processes in the ascending adjusted p value order. (C) Volcano plot depicting differentially regulated genes (black circles); red circles depict the significantly up- or down-regulated genes in *Plg^-/-^* compared with *Plg^+/+^*mice. (D) Heat maps showing significantly differentially expressed genes involved in granulopoiesis/neutrophil recruitment, myeloid cell activation and inflammation/extracellular matrix disassembly in *Plg^-/-^* compared with *Plg^+/+^* gingival tissue. (E) Flow cytometry analysis of 12- and 24-week-old mouse oral mucosal tissues. Contour plots show changes in neutrophil and monocyte/macrophage populations in percentages. Absolute count and percentages of (F) neutrophils and (G) monocytes/macrophages in 12- and 24-week-old *Plg^+/+^* and *Plg^-/-^* mouse gingiva. (H) Fibrin(ogen) and Ly6G IF staining of *Plg^-/-^* mouse oral sections showing localization of fibrin deposits and neutrophils (20x, Scale bar = 50 μm) and (I) Number of neutrophils/μm^2^ x 10^5^ in stained tissue sections.

To further characterize the lesion of immunopathology in *Plg^-/-^* mice, we next evaluated the gingival inflammatory infiltrate by flow cytometry (Gating strategy: Supplementary Figure 2). Consistent with the transcriptomics findings, this analysis revealed that the oral mucosal inflammatory infiltrate in *Plg^-/-^* mice is dominated by neutrophils (Live/CD45^+^/CD11b^+^/Cd11c^Neg-Med^/Ly6G^+^) at both early (12 weeks) and late (24 weeks) time points (Figure 2E and F). The absolute number of neutrophils was significantly increased in *Plg^-/-^* mice at all time points whereas their proportion increased significantly in late stages of immunopathology. In fact, neutrophils were the only cell subset significantly increased at early time points of immunopathology (12 weeks).

Monocyte/macrophage cells (Live/SSC^Int^/CD45^+^/CD11b^+^/Cd11c^Neg-Med^/Ly6G^-^) were not increased (neither in percentage or absolute numbers) at 12 weeks but displayed an increase in absolute counts at late stages of immunopathology (24 weeks), without a shift in proportion (Figure 2E and G). T cells (Live/CD45^+^/TCRβ^+^) and B cells (Live/CD45^+^/B220^+^) also did not show significant changes in proportion and numbers in early or late stages of immunopathology (Supplementary Figure 3).

Immuno-staining for fibrin(ogen) and Ly6G, a marker for neutrophils, in gingival tissues confirmed that neutrophils indeed accumulate in increased numbers in gingival tissues of *Plg^-/-^* mice and are in close proximity to fibrin(ogen) deposits (Figure 2H-I). Collectively, these data demonstrate that oral mucosal lesions in *Plg^-/-^* mice are dominated by neutrophils and suggest a neutrophil-mediated immunopathology.

### Fibrin(ogen)-mediated periodontal immunopathology depends on its myeloid integrin a.Mp2-binding motif

Conversion of soluble fibrinogen to the insoluble fibrin polymer exposes an otherwise concealed/cryptic binding motif for the integrin α_M_β_2_ that facilitates fibrin engagement with myeloid cells [16]. We hypothesized that neutrophil engagement of fibrin through α_M_β_2_ in gingival tissues is critical for neutrophil retention and/or activation and associated immunopathology. We tested this hypothesis by interbreeding *Plg^-/-^* mice with *Fgg^390-396A/390-396A^* mice, which bear a mutation in the concealed/cryptic binding motif on fibrin that mediates its engagement with myeloid cells. These mice have normal hemostasis, but they express a fibrin(ogen) that is unable to interact with α_M_β_2_ integrin [17]. The generated littermates were then assessed for periodontal bone loss. Interestingly, periodontal bone loss was completely prevented in *Plg^-/-^* mice by the elimination of the α_M_β_2_-binding motif from fibrin(ogen), as shown by comparing *Plg^-/-^*;*Fgg^390-396A/390-396A^* and *Plg^-/-^;Fgg^+/390-396A^* littermates (Figure 3A-B). Importantly, this prevention of periodontal bone loss in *Plg^-/-^* mice was not a consequence of diminished gingival fibrin accumulation, as fibrin(ogen) staining of the gingival tissues showed increased fibrin(ogen) accumulation in *Plg^-/-^*;*Fgg^+/390-396A^* and *Plg^-/-^;Fgg^390-396A/390-396A^* mice compared to *Plg^+/-^;Fgg^+/390-396A^* littermates (Figure 3C-D). More importantly, the prevention of alveolar bone loss in *Plg^-/-^* mice by elimination of the α_M_β_2_-binding motif on fibrin was also not a consequence of abolished gingival neutrophil accumulation. In fact, immuno-staining for tissue neutrophils (Ly6G) (Figure 3C and E), and multicolor flow cytometry of dissociated oral mucosal tissues demonstrated that *Plg^-/-^;Fgg^390-396A/390-396A^* mice have neutrophil accumulation comparable to *Plg^-/-^;Fgg^+/390-396A^* littermates (Figure 3F-G). Taken together, these data show that the local neutrophil engagement of fibrin(ogen) through α_M_β_2_ is essential for mucosal immunopathology and associated periodontal bone loss.

**Figure 3:**
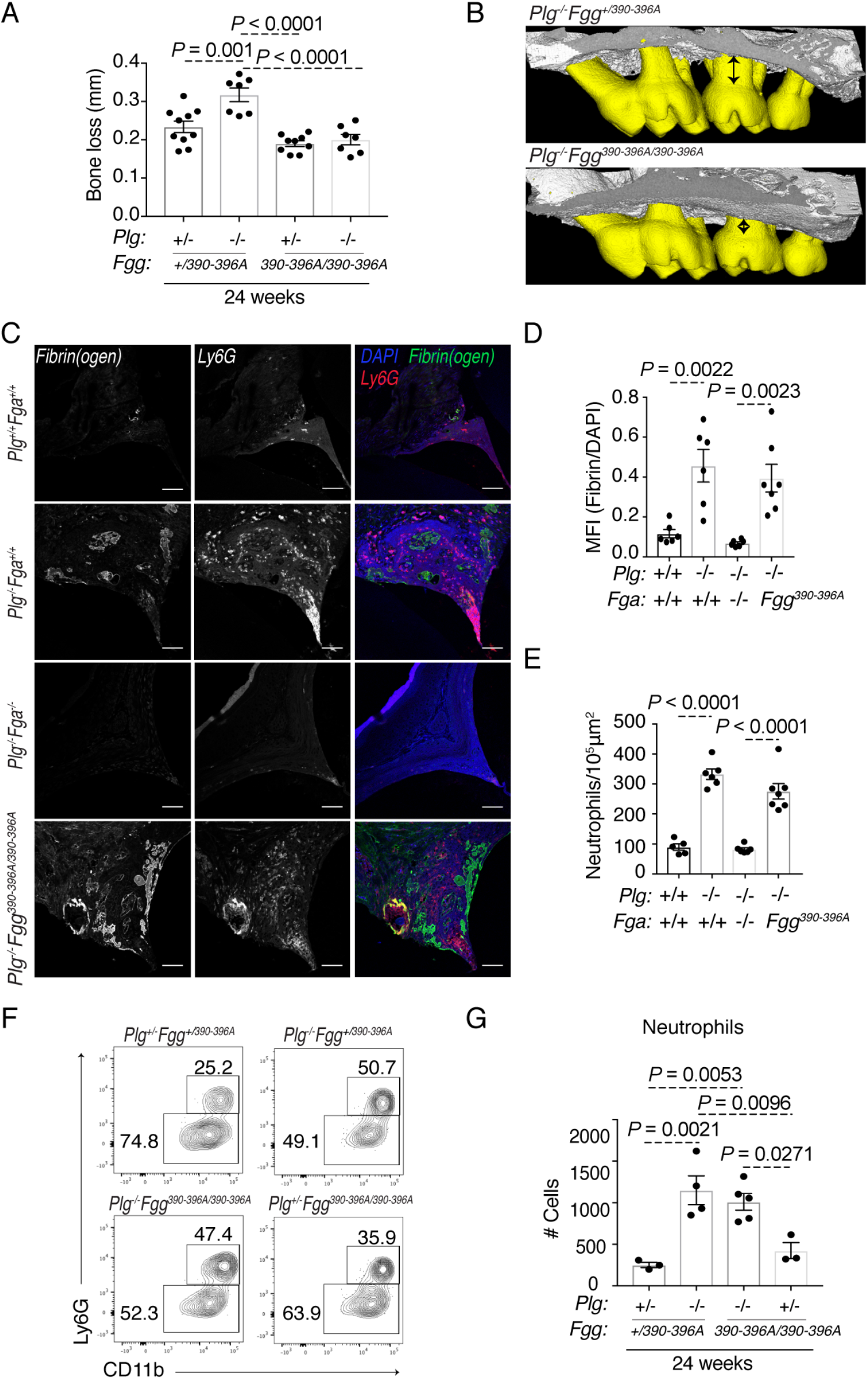
Myeloid integrin α_M_β_2_-binding motif on fibrin(ogen) mediates plasmin deficiency-induced oral immunopathology. (A) Bone loss measurement in *Plg^+/-^;Fgg^+/390-396A^, Plg^-/-^;Fgg^+/390-396A^, Plg^+/-^;Fgg^390-396A/390-396A^, Plg^-/-^;Fgg^390-396A/390-396A^* mice maxillae. (B) μCT visualization of periodontal bone loss of maxilla (black arrows depict the distance between alveolar bone crest and cemento-enamel junction). (C) IF staining of fibrin(ogen) and Ly6G in *Plg^+/-^;Fga^+/+^, Plg^-/-^;Fga^+/+^*, *Plg^-/-^;Fga^-/-^* and *Plg^-/-^;Fgg^390-396A/390-396A^* mouse gingival sections. Scale bar: 50 μm. (D) Mean fluorescence intensity of MFI (fibrin(ogen))/DAPI staining, and (E) number of neutrophils/μm^2^ x 10^5^ in stained tissue sections. (F) Flow cytometry analysis of 24-week-old oral tissue. Contour plots show neutrophil and monocyte/macrophage populations in percentages. (F) Counts of neutrophils in 24-week-old *Plg^+/-^*;*Fgg^+/390-396A^*, *Plg^-/-^;Fgg^+/390-396A^*, *Plg^+/-^*;*Fgg^390-396A/390-396A^*, and *Plg^-/-^*;*Fgg^390-396A/390-396A^* mouse gingival sections.

### Engagement of fibrin(ogen) through α_M_β_2_ integrins regulates neutrophil effector function

Next, we established an *in vitro* system to study the effects of engagement of neutrophils with fibrin(ogen) purified from either wild-type or *Fgg^390-396A/390-396A^* mice. Compatible with previous studies, human and mouse neutrophils showed a reduced adherence to mutant fibrin compared with wild-type fibrin (Figure 4A, Supplementary Figure 4 [18]). A similar reduction in adherence was noted when aM-deficient neutrophils were plated on wild-type fibrin and compared with wildtype neutrophils (Figure 4B), collectively affirming that α_M_β_2_ is important for neutrophil adhesion to fibrin. To assess the effect of neutrophil engagement of fibrin on neutrophil effector functions, we studied reactive oxygen species (ROS) production and neutrophil extracellular trap formation (NETosis) by human neutrophils plated on mutant or wild-type fibrin. We found that when neutrophils are in association with wild-type fibrin, ROS production per cell (Figure 4C-D) was significantly increased compared to neutrophils plated on fibrin lacking the α_M_β_2_ binding motif. Percentage of NETosis events as visualized by an increase in cellular size and subsequent externalization of DNA and myeloperoxidase (MPO) (Figure 4E-G, Supplemental Video and Supplementary Figure 5) were also significantly increased when neutrophils plated on wild-type fibrin were compared to fibrin lacking the α_M_β_2_ binding motif.

**Figure 4:**
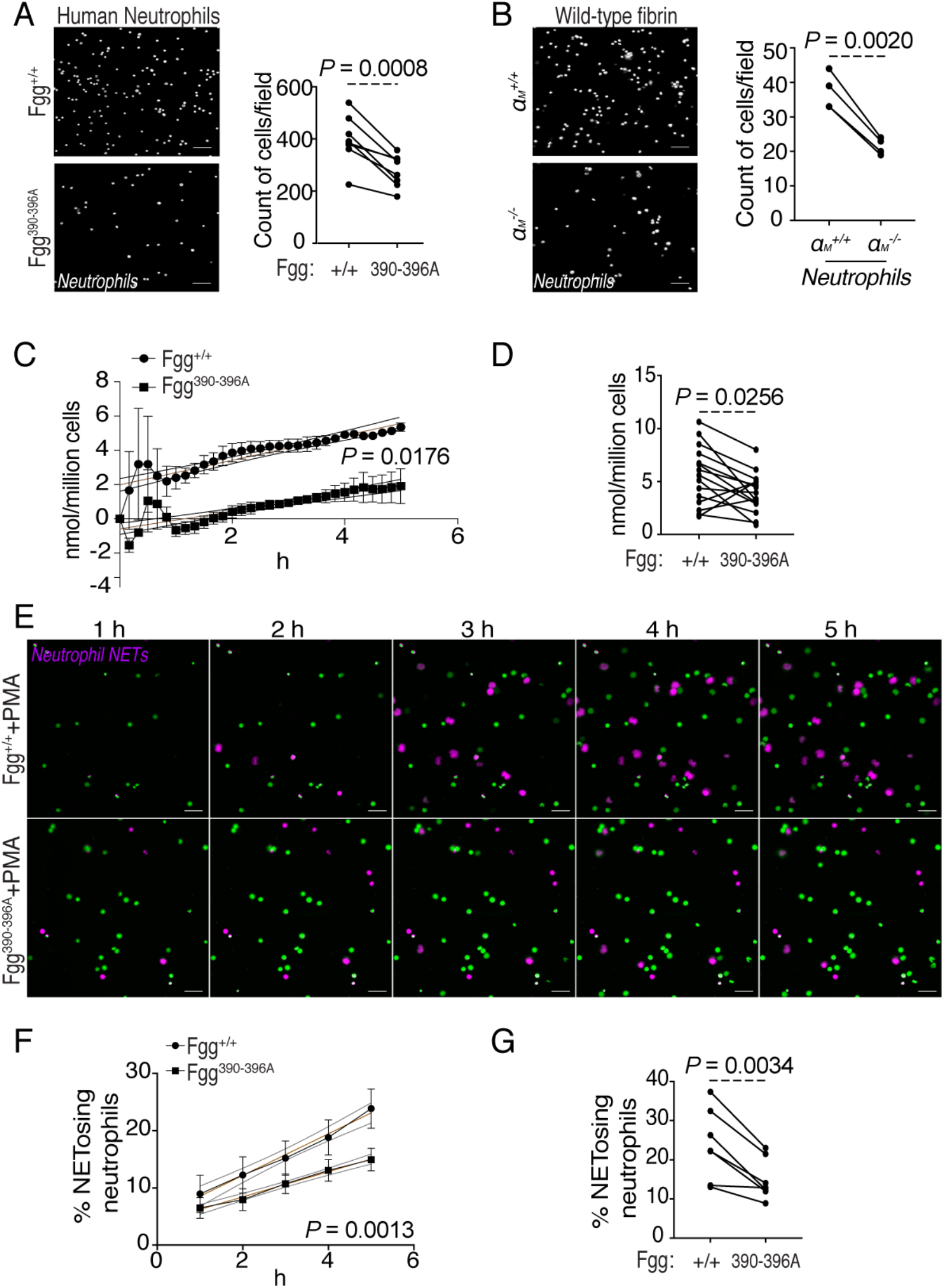
Fibrin(ogen)-neutrophil interaction through α_M_β_2_-binding triggers key neutrophil effector functions. (A) Human neutrophil binding to *Fgg^390-396A/390-396A^* fibrin compared to wildtype fibrin (40x; Scale bar = 200 μm). (B) Binding of wild-type and α_M_ integrin deficient (*α_M_^-/-^*) mouse neutrophils to wild-type fibrin (40x; Scale bar = 200 μm). (C-G) Neutrophils were plated on wild-type or Fgg^390-396A^ fibrin. (C) Neutrophil reactive oxygen species (ROS) production over time (15 nM MPO), and (D) ROS production at 5 h. (E) Neutrophil extracellular trap (NET) formation over time for neutrophils plated on wild-type and Fgg^390-396A^ fibrin (20x; Scale Bar = 50 μm), (F) and (G) Graphical representation of data depicted in panel E (Supplemental video).

Given the finding that fibrin regulates neutrophil effector functions through α_M_β_2_ *in vitro,* we next sought to investigate whether fibrin(ogen)-neutrophil interactions through the α_M_β_2_ motif would also mediate neutrophil activation *in vivo.* For this, we examined the spatial distribution of MPO in gingival tissues of *Plg^-/-^* mice expressing the mutated fibrinogen (*Plg^-/-^*;*Fgg^390-396A/390-396A^*) compared to those expressing wild-type fibrinogen (*Plg^-/-^;Fgg^+/+^*). Predicated on the assumption that MPO expelled by NETosing neutrophils, as supported by our *in vitro* studies, would appear more diffuse in tissue sections than MPO retained in granules of intact neutrophils, and therefore cover a larger area of a tissue section, we performed quantitative IF staining of gingival sections for Ly6G and MPO. *Plg^-/-^*;*Fgg^+/390-396A^* and *Plg^-/-^;Fgg^390-396A/390-396A^* mice gingival tissues showed comparable numbers of neutrophil accumulation. However, MPO staining of *Plg^-/-^*;*Fgg^+/390-396A^* mice covered a significantly larger area compared to *Plg^-/-^*;*Fgg^390-396A/390-396A^* littermates (Figure 5A-B), suggesting cellular externalization of MPO in *Plg^-/-^*;*Fgg^+/390-396A^* tissues consistent with NETosis (Figure 5A and C). We next aimed to investigate whether neutrophil NETosis is an important mechanism mediating immunopathology in the setting of Plg deficiency. For this, we treated *Plg^-/-^* mice with DNase I (i.p.) from week 8 to 20 and evaluated whether removal (or reduction) of neutrophil NETs (extracellular DNA) by DNase I treatment would reduce mucosal immunopathology. Importantly, removal (or reduction) of neutrophil NETs by systemic DNase I treatment (*i.p.*), significantly reduced alveolar bone loss in *Plg^-/-^* mice (Figure 5D-E), affirming that neutrophil NETosis *in vivo* is implicated as a mediator of oral mucosal immunopathology.

**Figure 5:**
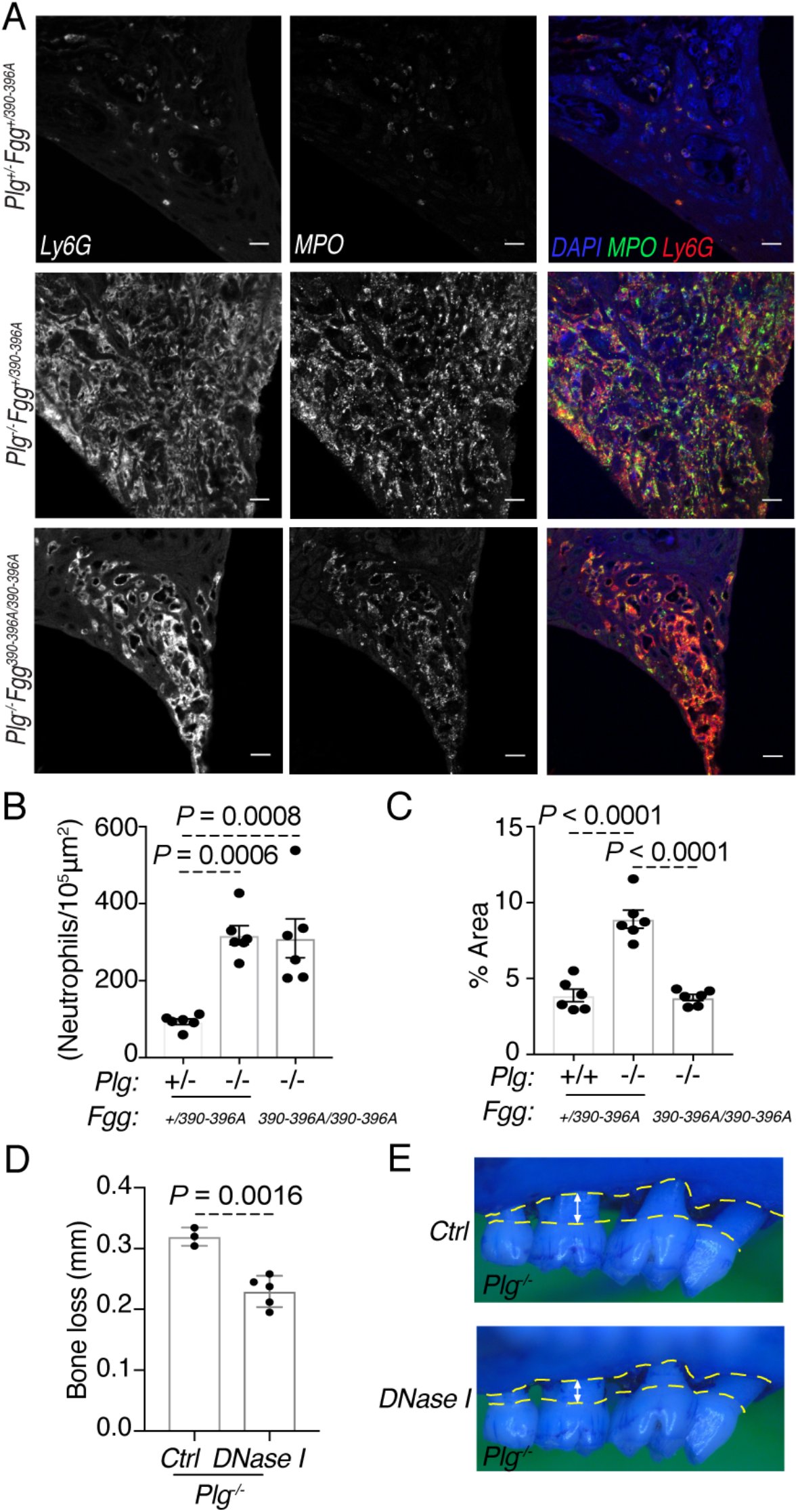
Neutrophil NETosis is a trigger for oral mucosal immunopathology in Plg-deficiency. (A) Immunofluorescence (IF) staining of Ly6G and MPO in *Plg^+/+^, Plg^-/-^* and *Plg^-/-;^Fgg^390-396A/390-396A^* mouse oral mucosal tissue sections (20x). Scale bar = 50 μm. (B) Number of neutrophils/10^5^ μm^2^ in stained tissue sections and (C) percentage of AI647 (MPO) stained area over total area. (D-E) *Plg^-/-^* mice were treated with DNase I (400 units twice a week *i.p*.) or vehicle control (0.9% NaCl). Bone loss measurement after DNase I or vehicle treatment.

### Fibrin(ogen) engagement with the β_2_ integrin promotes oral mucosal immunopathology in the context of normal fibrinolysis

Our work reveals fibrin(ogen)-mediated neutrophil activation as an essential pathway in oral mucosal immunopathology, particularly in the context of Plg deficiency, where excessive extravascular fibrin deposition is observed. However, we theorize in the setting of periodontitis, perpetual microbial stimulation may also lead to increased fibrin(ogen) accumulation, neutrophil recruitment and subsequent fibrin-neutrophil-mediated immunopathology. To address this hypothesis, we explored a model of age-dependent periodontitis in mice [19]. Wild-type mice develop periodontal bone loss with age. By 24 weeks of age, wild-type mice will typically display significant bone loss compared to younger (8-10 week old mice) ([20] and Fig 1A). Therefore, we interrogated whether fibrin(ogen)-mediated neutrophil activation may play a role in the natural progression of periodontitis with age. For this, we aged *Fgg^390-396A/390-396A^* mice to 24 weeks and evaluated periodontal bone loss compared to littermate controls. Interestingly, *Fgg^390-396A/390-396A^* mice had significantly reduced periodontal bone loss compared with wild-type and heterozygous littermate controls, demonstrating that the fibrin(ogen) α_M_β_2_ binding motif is critical for the development of periodontal bone loss even under normal fibrinolytic conditions (Figure 6A).

**Figure 6:**
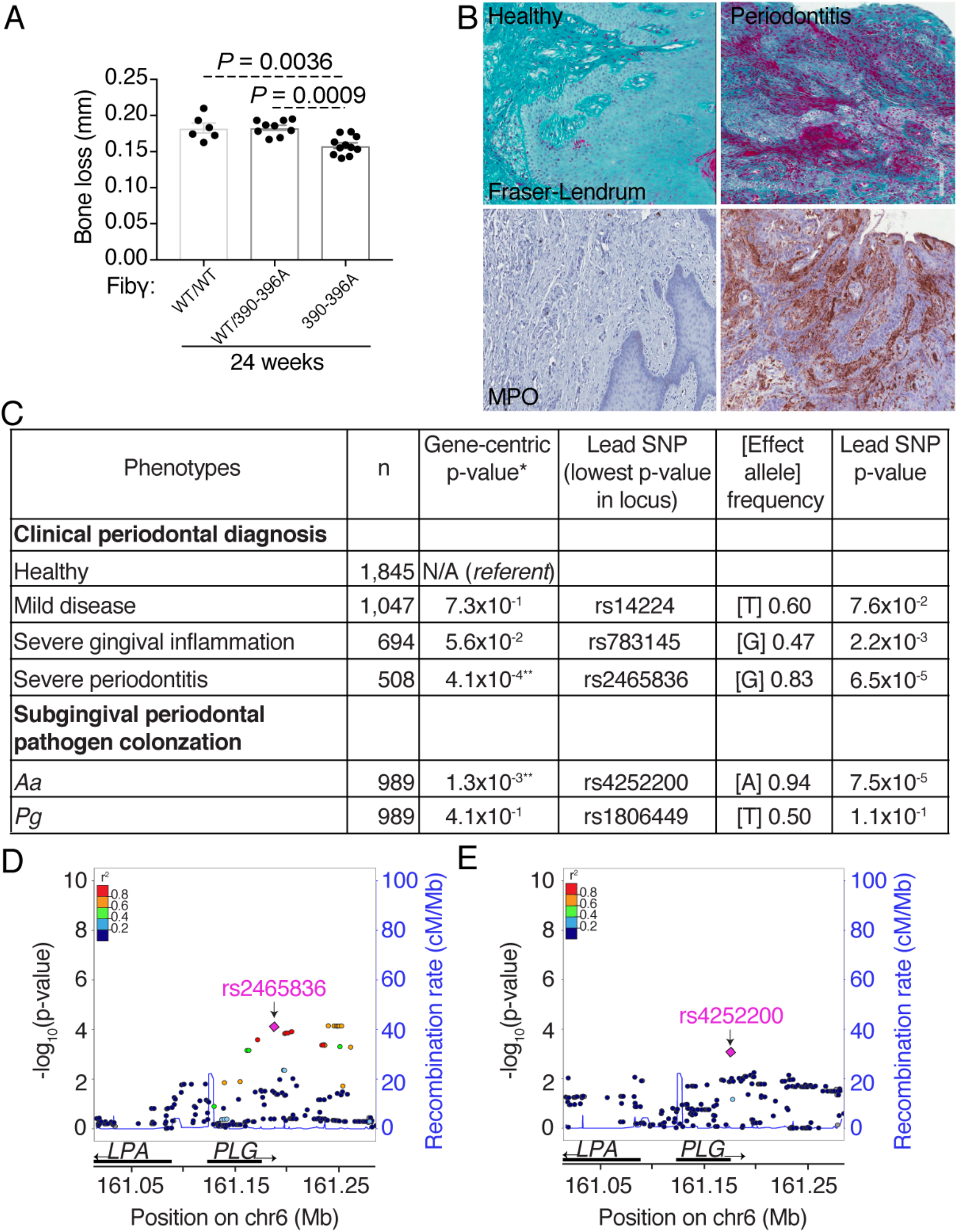
Fibrin-neutrophil axis as an initiator in common forms of periodontitis. (A) Bone loss measurement in *Fgg^+/+^, Fgg^+/390-396A^* and *Fgg^390-396A/390-396A^* mice maxillae at 24 weeks of age. (B) Fraser-Lendrum staining for fibrin(ogen) and immunohistochemistry (IHC) staining of MPO in oral mucosal tissues from patients with severe periodontitis and healthy controls. (C) Results of gene-centric association testing of *PLG* polymorphism with clinical and microbiologic data. *denotes gene-centric association p-values obtained using MAGENTA (using a 110Kb upstream

These experimental data in mice led us to consider whether the fibrin-neutrophil axis could be implicated in common (non-Mendelian) forms of human periodontitis. In periodontitis, local tissue inflammation and bleeding is a hallmark of disease [21]. Additionally, infiltration of neutrophils is a characteristic of periodontitis [22]. Consistent with the notion that neutrophil-fibrin(ogen) interactions could potentially also mediate periodontitis in patients, we found accumulation of extravascular matrix protein (consistent with fibrin, Fig. 6B) in lesions of human periodontitis but not in healthy tissues. Furthermore, robust neutrophil infiltrates were observed in these same tissue lesions of periodontitis patients (Fig. 6B).

### *PLG* polymorphisms are associated with the common oral mucosal disease periodontitis

Genomic data support the role of Plg-dependent pathways in common forms of periodontitis. Recent genome-wide association studies (GWAS) reported that polymorphisms downstream of *PLG* (rs1247559) were associated with aggressive periodontitis in Europeans [23, 24]. In agreement with these previous reports, we generated new evidence of *PLG* association with clinical and microbiological parameters of periodontal disease in a cohort of adults of European ancestry in North America. We found that *PLG* polymorphisms *in tandem* are associated with severe periodontitis (*PLG* gene-centric p=4.2×10^−4^; lead SNP: rs2465836) and with high colonization of the periodontal microbe *Aggregatibacter actinomycetemcomitans (Aa),* which is associated with aggressive periodontitis (*Aa* colonization, gene-centric p=1.3×10^−3^; lead SNP: rs4252200) (Fig. 6C-E). Mild periodontal disease, gingival inflammation (without periodontal destruction, Supplementary Figure 7) and colonization with the microbe *Porphyromonas gingivalis (Pg,* which is associated with chronic forms of periodontitis), did not yield any material evidence of association with *PLG* polymorphisms (Fig. 6C-E). These new, independent findings are consistent with previous reports of periodontitis risk-conferring signals in the region immediately downstream of *PLG* [23] [24] (Fig. 6D-E). These findings together with previous reports provide a summary of evidence linking genetic variation in the *PLG* locus with severe and potentially aggressive forms of human periodontitis.

## Discussion

In this study, we show that plasmin-dependent fibrinolysis is essential for mucosal barrier homeostasis. We further demonstrate that accumulation of fibrin is a trigger for mucosal immunopathology through local activation of neutrophil effector functions.

We document, consistent with a previous study, that *Plg^-/-^* mice develop severe periodontitis [14]. We confirm that this disease is mediated by the enzymatic activity of plasmin, as mice with either (i) genetic ablation of Plg activators *Plat^-/-^*;*Plau^-/-^* or (ii) a mutation in the catalytic serine residue of plasmin, which renders plasmin catalytically inactive (*Plg^S743A/S743A^*), both display spontaneous oral mucosal destruction. We demonstrate that the critical substrate of plasmin, which ultimately mediates disease, is fibrin(ogen). Genetic elimination of fibrin(ogen) from mice lacking Plg (*Plg^-/-^;Fga^-/-^*) completely prevented the Plg deficiency-associated periodontal bone loss, revealing that oral mucosal immunopathology in Plg deficiency is mediated by fibrin(ogen).

Fibrin(ogen) harbors a cryptic β_2_ integrin-binding motif, which is specific for the β2 integrins on myeloid cells, and becomes exposed upon conversion of fibrinogen to fibrin [25, 26]. Mutations in the β_2_ binding motif protects from fibrin(ogen)-associated immunopathology in models of autoimmune disease, including arthritis and multiple sclerosis, due to disruption of monocyte/macrophage/microglial activation [27–29]. However, mechanisms underlying fibrin-triggered mucosal disease and fibrin-neutrophil mediated immunopathology are not well understood. We demonstrate that mucosal immunopathology is mediated through the β_2_ binding motif on fibrin. Expression of *Fgg^390-396A^* fibrin(ogen) which lacks the integrin binding motif, completely prevented periodontitis in *Plg^-/-^* mice. Importantly, while we observed a complete prevention of periodontal bone loss in *Plg^-/-^* mice expressing *Fgg^390-396A^* fibrin(ogen) (*Plg^-/-^*;*Fgg^390-396A^*), this was not due to either reduced fibrin(ogen) or reduced neutrophil accumulation. In fact, we observed a similar level of fibrin(ogen) and neutrophils in both *Plg^-/-^;Fgg^390-396A^* mice and *Plg^-/-^* mice gingival lesions. We therefore concluded that fibrin mediates immunopathology through a “functional engagement” of local neutrophils.

We propose that fibrin-mediated neutrophil activation is an essential mechanism for mucosal homeostasis. In our *in vitro* model system, we confirm that neutrophil adherence to fibrin is mediated by both the α_M_β_2_ binding motif on fibrin and by the expression of the α_M_ integrin subunit on neutrophils. We show that fibrin(ogen)-neutrophil engagement is essential for the execution of key neutrophil effector functions, including reactive oxygen species (ROS) production, myeloperoxidase expression and neutrophil extracellular trap formation (NETosis) [30]. Fibrin-neutrophil engagement through the β_2_ binding motif requires exposure of the cryptic binding domain on fibrin. Exposure of this motif occurs wither when fibrinogen is converted to fibrin. Thus, we suggest that this context-dependent mechanism of fibrin(ogen) ligand exposure is an important pathway by which neutrophils become activated within tissue microenvironments. At barrier and mucosal sites, constant environmental triggering leads to perpetual inflammatory stimulation. Deposition of extravascular fibrin through activation of the coagulation cascade is a universal aspect of inflammation. Neutrophils arrive at sites of injury as part of the early immune response, which is geared towards anti-microbial defense and wound clearance. We thus theorize that neutrophil activation in a wound bed may be mediated through the fibrin matrix in order to trigger local bacterial clearance. Indeed, NET formation is considered a major mechanism for neutrophil-mediated containment of microbes [31], particularly those that are too large for phagocytosis [32]. However, in the setting of genetic defects in fibrinolysis, excessive mucosal fibrin deposition will perpetually engage and activate neutrophils leading to local immunopathology. In this scenario, prolonged and/or exaggerated activation of neutrophils will trigger production of ROS and release of NETs, and has been linked to a variety of inflammatory and autoimmune pathologies [33–35].

Release of NETs has been shown to amplify immune responses in several diseases [36, 37]. In the setting of periodontitis, both release of ROS and NETs can contribute to tissue destruction and bone loss. Excessive ROS can exert direct cytotoxic effects on matrix fibroblasts [38], but has also been reported to be involved in periodontal bone destruction through the stimulation of osteoclast formation [39, 40]. Furthermore, release of NETs has been shown to induce bone loss in models of arthritis [41], particularly through activation of resident tissue fibroblasts [42]. Diverse stimuli have been shown to induce NET release [43], with ROS-and MPO-dependent release of NETs being one of the major pathways involved in NET release [36]. Importantly, consistent with our findings, engagement of β2 integrin on neutrophils has been previously shown to mediate ROS induction [44] and NET release in response to fungal stimuli [45].

Beyond discovery of critical mechanisms for mucosal immunity, our work provides valuable insights towards the understanding and therapeutic targeting of the oral mucosal disease periodontitis. This is a very common human condition, which affects in its moderate-severe forms 8-10% of the general population [46]. It is thought that, in periodontitis, microbial stimulation leads to excessive mucosal inflammation and consequent loss of bone. Mechanisms of disease are not completely dissected, and treatment largely depends on removal of the microbial biofilm rather than host targeting. The genetic basis of disease continues to be explored. Notably, genetic forms of periodontitis are largely seen in patients with dysfunctions in neutrophils [47, 48], suggesting neutrophil function as an important mediator of periodontal homeostasis [49]. In common forms of the disease, excessive neutrophil accumulation and activation, including NET formation has been documented [50–53]. Recently, neutrophil recruitment through induction of Th17 responses has been mechanistically linked to the pathogenesis of periodontitis [54]. However, how neutrophil activation is locally mediated and may participate in disease progression is not completely dissected. Based on our findings, we theorize that local accumulation of fibrin (secondary to microbiome-induced inflammation) is a mediator in common forms of periodontitis. Indeed, we document abundant fibrin deposition and neutrophil accumulation in patient lesions of periodontitis. We also provide mechanistic evidence that fibrin-mediated inflammation, specifically engagement of the α_M_β_2_ binding motif, can contribute to periodontal bone loss in animal models even in the absence of fibrinolytic deficits. We show that mice with the *Fgg^390-396A/390-396A^* mutation in the α_M_β_2_ binding site on fibrin were protected from spontaneous age-related periodontal disease, indicating that activation of neutrophil effector functions through fibrin-engagement contributes to periodontal disease in the setting of both compromised and normal fibrinolytic function. Importantly, this pathway of fibrin-mediated inflammation could be therapeutically targeted in humans. Recently, a monoclonal antibody has been generated to target the cryptic α_M_β_2_-binding fibrin epitope γ377–395, and selectively inhibit fibrin-induced inflammation in neurological disease [55]. Conceivably, this therapeutic modality would be applicable for inhibition of inflammation in periodontitis.

Notably, human genetic evidence supports the role of *PLG* in common forms of periodontitis, in the absence of a Mendelian disease. A recent meta-analysis of genome-wide association studies of periodontitis comprising almost 15,000 adult participants of European ancestry, revealed *PLG* polymorphisms as one of the 2 major risk factors for aggressive periodontitis [23]. An earlier report has also highlighted polymorphisms downstream of *PLG* as a shared genetic risk locus between cardiovascular disease and periodontitis [24]. In agreement with these previous reports, we have now independently confirmed that *PLG* is a genetic risk factor for periodontitis in a separate cohort of adults of European Ancestry in North America. We document that polymorphisms downstream of *PLG* are associated with severe forms of periodontal disease. We also find a strong association of *PLG* polymorphisms with high levels of the periodontal microbe *Aggregatibacter actinomycetemcomitans (Aa),* a microbe typically detected in patients with aggressive forms of periodontal disease [56]. Interestingly, the lead SNP in our analysis associated with severe periodontitis (rs2465836) is in strong linkage disequilibrium with the lead marker (rs1247559) in the recent European report of aggressive periodontitis [23]. Moreover, the SNP (rs4252200) leading the signal for high *Aa* colonization is proximal to the previously reported aggressive periodontitis and cardiovascular disease-related polymorphism (rs4252120) and has also been reported as strongly associated with severe asthma triggered by fungal sensitization [57].

Collectively, our data reveal that fibrin operates as a critical immune regulator of mucosal barrier homeostasis and a potential pathogenic driver of mucosal disease through the regulation of neutrophil activation.

## Methods

### Mice

*Plg^-/-^*, *Plg^S743A/S743A^*, *Plat^-/-^;Plau^-/-^*, *α_M_^-/-^*, *Fga^-/-^* and *Fgg^390-396A/390-396A^* knock-in mice have been previously described [12, 13, 18, 58–61]. All studies were littermate controlled. Mice were in a C57BL/6J background, except for *Plg^-/-^*;*Fga^-/-^* mice and their *Plg^-/-^*;*Fga^+/-^*, *Plg^+/-^*;*Fga^-/-^* and *Plg^+/-^*;*Fga^+/-^* littermates, which were in the mixed Black-Swiss/C57BL/6J background. Mice were genotyped as previously described [12, 13, 18, 58–62]. All experiments were performed under approved protocols in Association for Assessment and Accreditation of Laboratory Animal Care International-certified vivarium and were approved by the Institutional Animal Care and Use Committee prior to initiation of experimentation.

### Bone loss measurements

Periodontal bone heights were assessed after defleshing and staining with methylene blue. The distance between the cemento-enamel junction and alveolar bone crest (CEJ-ABC distance) was measured at six predetermined sites as previously described [19] and combined to give a total CEJ-ABC distance. Results were displayed as mean±SEM.

### Micro-computed tomography (μ-CT)

Mouse maxillae were dissected at 24 weeks (6 mice for each age group), for quantitative analysis of the alveolar bone, hemi-maxillae were examined by a micro-CT system (μCT 50, Scanco Medical AG, Bassersdorf, Switzerland) as previously described [63]. Briefly, the sagittal plane of the specimens was set parallel to the x-ray beam axis. The specimens were scanned at a resolution of 4.4 μm in all three spatial dimensions at 90KV, 88μA, 600 seconds integration time. Reconstructed scans were then exported to AnalyzePro (AnalyzeDirect, vers. 1.0) to quantify volumetric changes in alveolar bone, among the groups of mice at 24 weeks.

#### Linear measurements

Hemi maxillae were reoriented to align the occlusal plane in all samples using AnalyzePro. All images were reoriented such that the CEJ and the root apex (RA) appeared in the micro-CT slice that was to be analyzed. Linear measurements were taken (in millimeters) from the cemento-enamel junction to the alveolar bone crest, similar to the periodontal bone height measurement above.

#### Volumetric micro-CT measurements

Volumetric measurements were carried out following the selection of a 3-D region of interest (ROI). The most mesial root of first molar and the most distal root of third molar served as endpoint borders because they were the most consistent among specimens. The height of the ROI was fixed at 0.7 mm from the most coronal point in the bifurcation of the 1^st^ and 3^rd^ molars (approx. 150 slices) in all samples. The resulting ROI was segmented by using the same threshold in all samples and the volume of the segmented region was calculated in AnalyzePro. The values were expressed as mean±SD.

### Fraser-Lendrum staining

Decalcification, paraffin embedding and staining of 5 μm sections of mouse maxillae and mandibles using the Fraser-Lendrum method was performed by Histoserv Inc. (Germantown, MD) [64].

### Immunofluorescence staining and quantification

Immunofluorescence confocal microscopy was performed using 5 μm sections of decalcified and paraffin embedded maxillae and mandibles (Histoserv Inc.). Antigen retrieval in 10 mM citric acid buffer was used, and sections were blocked with 2% bovine serum albumin in phosphate buffered saline (PBS) for 1 h. Sections were incubated with primary antibodies at 1:500 dilution [anti-fibrin(ogen): [58]; anti-Ly6G: BD Pharmingen™ 551459; anti-MPO: ab65871] O/N at 4 °C and washed in PBS (3 times). Secondary antibody (Rhodamine Red-X anti-rat, JacksonImmuno 712-296-150; Alexa Fluor® 647-conjugated AffiniPure F(ab’)2 anti-rabbit, JacksonImmuno 711-606-152; 1:500) and DAPI (1:1000) were added for 1 h at room temperature. Sections were then washed in PBS (3 times), and #1.5 cover slips were mounted with ProLong™ Gold Antifade Reagent (Thermo Fisher Scientific). Images were captured on an inverted Nikon A1R+ confocal microscope (20x-air or 60x-oil immersion) using NIS-Elements software (Nikon).

For fibrin quantification, the average fluorescent intensity was measured using NIS-Elements software. A fixed-area square region of interest was drawn around the tissue and the Time Measurement analysis tool was used to measure the average fluorescent intensity within the ROI for the AI555 channel. Five individual fields were measured per tissue and the mean fluorescent intensity values were calculated in Microsoft® Excel for Mac (version 16.15). Results were displayed as mean±SEM.

To quantify the myeloperoxidase-stained area in the gingiva, a statistical threshold (set at 2 standard deviations above the mean of the image histogram) was set. An investigator-drawn region of interest determined the appropriate area of the sample and the square area was reported. A similar analysis was applied to immunofluorescent images of Ly6G to determine the number of invading neutrophils to the gingival lesion. The area was calculated and divided by the average area of a neutrophil (determined here to be approximately 22.1 μm^2^, n=95) to estimate the number of invading neutrophils. The analysis was performed with MetaMorph and results were displayed as mean±SEM.

### RNA-Seq

Total RNA was extracted from experimental tissue samples and sequencing libraries were prepared by the Illumina Nextera XT method following manufacturer’s recommendations (Illumina, San Diego, CA). The multiplexed libraries were sequenced in a NextSeq500 instrument in 150 bp paired end mode. Demultiplexed samples were mapped and transcripts quantified to the GRCm38.v11 mouse genome using the STAR v2.5.2 aligner. Gene-level counts were filtered to remove low expression genes (keeping genes with > 5 counts in at least one sample). The filtered gene expression matrix was analyzed by principal component analysis to identify outlier samples (none detected). Differential gene expression was evaluated by three independent statistical methods (DESeq2, edgeR, and Limma-Voom). Differentially expressed genes (FDR < 0.05) were considered for further analyses, based on results from DESeq2.

### Preparation of single cell suspensions

Mice gingiva were dissected and digested for 50 min at 37 °C with Collagenase IV (GIBCO) and DNase (Sigma) as previously described [65].

### Flow cytometry

Single cell suspensions of gingiva were stained with the Live/Dead Cell Viability assay (Invitrogen), and cell surface markers were stained with the following anti-mouse antibodies at 1:200 dilution per 10^6^ cells: Ly6C (AL-21; BD Biosciences), Ly6G (1A8; BD Biosciences), B220 (RA3-6B2; BioLegend), CD11B (M1/70; BioLegend), TCRβ (H57-597; BioLegend), MHCII (M5/11.4.15.2; eBioscience), F4/80 (BM8; eBioscience), CD11C (N418; eBioscience) and CD45 (30.F11; eBioscience). All samples were analyzed using a FACS Fortessa cytometer (BD Biosciences). Data analysis was performed using FlowJo software (Tree Star). Results were displayed as mean±SEM.

### Purification of plasma fibrinogen

Wild-type and *Fgg^390-396A/390-396A^* murine fibrinogen was purified from citrated-plasma by ammonium sulfate precipitation as described [66].

### Neutrophil isolation from mouse bone marrow

Neutrophils were isolated as described previously [67]. Briefly, 8-to 12-week-old mouse bone marrow cells were flushed out from femurs and tibia with RPMI 1640 supplemented with 2 mM EDTA, 20% FBS and 1% Pen/Strep. Neutrophils were isolated from total bone marrow cells by density gradient centrifugation on Histopaque 1119/Histopaque 1077. The purity of the neutrophils was above 95% as determined by flow cytometry analysis. Then neutrophils were washed with PBS and suspended in RPMI 1640 supplemented with 2% heat-inactivated fetal bovine serum (FBS).

### Neutrophil isolation from human blood

Human neutrophils were isolated from heparinized peripheral blood from the NIH blood bank using density gradients of Ficoll-Paque premium (GE Healthcare, Pittsburg, PA) followed by removal of red blood cells (RBC) using dextran-promoted rosette formation and hypotonic lysis of remaining RBC.

### Cell-based assays

#### Fibrin coating of wells

Plates were coated with purified wild-type or mutated fibrin (0.60 μg/mL mouse fibrinogen + 5 U/mL mouse thrombin) at 37 °C O/N in a humidified incubator. Excess thrombin was washed off with DMEM and non-specific binding sites were blocked using 1% nonfat dry milk (1 h) followed by DMEM washing (3x). Plates were allowed to dry for 15 min before use.

### Cell adhesion assay

Cell adhesion assays were performed in 96 well Black μ-Plates (ibidi). Aliquots of cells (1×10^5^ cells/mL) were added to fibrin-coated wells, incubated at 37 °C for 30 min, and subsequently washed with buffered saline to remove non-adherent cells. The number of adherent cells was determined by counting DAPI-stained cells in multiple high-powered (40x) fields. All analyses were performed in triplicate.

### Reactive oxygen species (ROS) production

ROS assay was performed in 96 well clear plates (ThermoFisher Scientific). Aliquots (100 μL) of cells (3.3×10^6^ cells/mL) were added to wells and incubated at 37 °C for 30 min. Control wells without cells were used to negate any contaminant from fibrin. Cells were supplemented with (25 μL) cytochrome-c (1800 nM) with or without 15 nM PMA (phorbol 12-myristate 13-acetate) and/or superoxide dismutase (SOD; 60 U/mL) and absorbance was measured at 550 nm for 5 h every 10 min. The absorbance values at 550 nm (differences in OD between samples with and without SOD) were converted to nanomoles of O_2_-based on the extinction coefficient of cytochrome-c: D_E550_ = 21 × 10^3^ M^-1^cm^-1^. The results were expressed as nanomoles of O_2_-per 1×10^6^ cells.

### Neutrophil extracellular trap (NET) formation

NET formation assays were performed in 8 well μ-Slides (ibidi). Aliquots (200 μL) of cells (2.5×10^4^ cells/mL) and Hoechst (1 μg/mL) were added per well and incubated at 37 °C for 30 min in a humidified incubator. Wells were washed with FluoroBrite DMEM (Thermo Fisher Scientific) to remove non-adherent cells and supplemented with 200 μL of FluoroBrite DMEM containing 0.2 μM Sytox Green. Time-lapse series were acquired every 5 min for 5 h on an inverted Nikon TI-E microscope using a Plan Fluor 20x (N.A. 0.5) objective and a Flash-4 v3 sCMOS camera (Hamamatsu). Excitation and emission were provided by a Sola light engine (Lumencor) and standard DAPI and GFP filter sets (Chroma) for Hoechst and Sytox Green, respectively. Light intensity and camera exposure times varied between 3-5% and 100 ms, respectively. An environmental chamber (Precision Plastics) kept samples humidified and at a constant 37 °C with 5% CO2. All hardware was controlled through Nikon Elements. After the first 10 min of imaging, cells were supplemented with 200 μL of FluoroBrite DMEM containing 0.2 μM Sytox Green with or without 15 nM PMA and continued imaging for 5 h.

### Neutrophil extracellular trap (NET) analysis and quantification

After live cell imaging, all time-lapse files were corrected for drift using the “Correct 3D drift” plugin in Fiji (ImageJ 2). For an example of the processing and analysis, see Supplementary Figure 6. To distinguish cells undergoing apoptosis from NETosis, we generated Fiji scripts in the following manner: The final image of the Cytox Green channel was duplicated and subtracted from the entire time series using the postulation that any cell remaining green after 5 hrs was apoptotic and had not fully ruptured or undergone chromosome decondensation. This gave us a “NETosis only” image stack. This “NETosis only” stack was then subtracted from the original to create a “Apoptosis only” stack. Substacks of each were created that represented 1-hr, 2-hr, 3-hr, 4-hr, and 5-hr. No image convolution filters were used for apoptosis or NETosis data. Maximum intensity projection (MIP) renderings were generated from these substacks and then concatenated. An Otsu threshold was applied to each image stack and measured using the “Analyze Particles” plugin using different size filters to exclude smaller cells (NETosis>120; Apoptosis>52). For some samples where an automated threshold could not be used, the user visually set the threshold to properly cover the cell edge. The Hoechst channel was used to determine live from dead cells. An unsharpen mask (radius=15, 0.60 scaling) was used to remove out of focus blur. Otsu Thresholding was then applied and the “Analyze Particles” plugin was again used to exclude random noise (>10). The user then visually determined if each was correct and adjusted false positives prior to any further statistical analysis. Please contact the authors for the actual scripts or see the authors’ github at https://github.com/addoyle1D/NETosis.

### DNase I treatment

*Plg^-/-^* mice of 12 weeks of age were *i.p.* injected with either 400 units of DNase I (Millipore Sigma) in 100 uL of 0.9% NaCl or 0.9% NaCl, twice a week for 8 weeks. Heads were harvested and bone loss measurements were taken as described under “Bone loss measurements”.

### Human oral mucosal samples

Biopsy samples form oral mucosal tissues surrounding teeth (gingiva) were harvested from healthy volunteers and patients with severe-untreated periodontal disease. All patients were enrolled in an IRB-approved clinical study at the NIH hospital (NCT#01568697) and provided informed consent. Human biopsies were stained for MPO (myeloperoxidase) as follows and fibrin (Fraser Lendrum stain) as established before. Antigen retrieval in 10 mM citric acid buffer was used, and sections were blocked with 5% bovine serum albumin in phosphate buffered saline (PBS) for 1 h. Sections were incubated with MPO (ab9535) at 1:500 dilution. Secondary antibody anti-rabbit (ImmPRESS HRP Reagent kit, Vector Laboratories) for 30 minutes, followed by washing three times in PBS then adding developing solution (ImmPACT DAB, Vector Laboratories). The reaction stopped with water and slides counterstained with Hematoxylin QS (Vector Laboratories).

### Human genetic associations with clinical and microbiological periodontal disease parameters

To examine the association of polymorphisms in the *PLG* gene locus with the presence and severity of human periodontal disease in an independent cohort, we used existing clinical, microbiological and genotype information from the Atherosclerosis Risk In Communities (ARIC) study [68]. ARIC is a prospective cohort study of atherosclerosis and cardiovascular disease risk factors which included a Dental ARIC ancillary study with comprehensive oral examinations in a subset of ARIC participants [n=4,504 European Americans (EA)]. Oral microbiome samples were collected from subgingival tooth surfaces in a subset of Dental ARIC participants (n=1,020 EA) and were subsequently analyzed for the presence/abundance of eight common periodontal pathogens. Oral disease diagnoses were based on the recently introduced P^3^ classification system [69], separating the population into 7 distinct oral health categories. Four categories focused on the severity of periodontal disease [mild disease, severe inflammation (inflammation without periodontal bone loss), localized disease and severe periodontitis (severe periodontal bone loss)]. Two categories reflected oral disease burden not specific to periodontitis (tooth loss and severe tooth loss) and the final category is designated for periodontal health [70]. Microbiome-specific categories were assigned for patients with high detection of either *Porphyromonas gingivalis* (*Pg*, associated with the most common, chronic types of periodontitis) [71] or *Aggregatibacter actinomycetemcomitans* (*Aa*, associated with aggressive periodontitis) [56].

We used existing genotype data that were initially generated using the Affymetrix Genome-Wide Human SNP Array 6.0 chip [n~900,000 markers for single nucleotide polymorphisms (SNPs)] and then imputed to the HapMap Phase II CEU build 36 panel resulting to a final analysis set of 2.1 million SNPs with minor allele frequency of >5% [68]. Genetic association models for single markers (i.e., SNPs) were based on logistic regression (i.e., “disease” vs. “healthy” periodontal groups and “high” vs. “lower” pathogen colonization), assuming log-additive allelic effects, and including adjustment terms of study design, 10 principal components for ancestry, age and sex. To evaluate *PLG* genetic association with clinical and microbiological parameters of periodontal disease first we defined an extended gene region around it, 110Kb upstream to 40Kb downstream of the gene boundaries, harboring 153 SNPs. The selection of the specific boundaries was based upon simulation and actual data applications of MAGENTA [72] that was used to obtain genecentric association estimates. We used the MAGENTA approach to synthesize single-marker association signals in the defined *PLG* gene region into a single association estimate for *PLG* (Z score and corresponding p-value), accounting for the number of SNPs tested and their local linkage disequilibrium (LD) structure, based on the human genome build 36. Here, we obtained *PLG-* centric estimates of association with clinical diagnoses and microbiological periodontal parameters and report gene p-values, as well as information for the lead (i.e., lowest p-value) SNP in the gene region. In this study, we formally tested for 5 *PLG*-trait associations (“mild periodontitis”, “severe gingival inflammation”, “severe periodontitis”, “high” *Aa* and “high” *Pg* colonization). Because the P^3^ system includes 6 clinical disease groups (including localized disease and patterns of tooth loss), and since 4 periodontal pathogen colonization traits have been previously reported, we applied a conservative Bonferroni multiple testing correction assuming 10 independent tests (critical value p<5×10^−3^). Finally, to illustrate and provide more information about the genomic context of the identified *PLG* locus associations, we generated regional association (Locus Zoom) plots [73] using single-marker association results in areas ±110 Kb flanking *PLG.*

### Statistical Analysis

For *in vivo* assays, the unpaired Student’s t-test or the one-way ANOVA (when >2 groups), unpaired two-tailed statistical analysis was performed. For *in vitro* assays, results were displayed as mean±SEM and the two-tailed, Wilcoxon signed-rank test was performed. For time point analyses (ROS and NETosis) linear regression analysis was performed to test whether differences in slopes were significantly different. 95% confidence interval lines were displayed on graph. All statistical analyses were performed on Graph Pad Prism (v7).

## Acknowledgements

We thank Dr. Mary Jo Danton and Dr. M Lionakis for critically reviewing this manuscript. This research was funded by the NIDCR Intramural Research Program (T.H.B, and N.M.M), and supported by the NIDCR Veterinary Resources Core (ZIC DE000740-05), the NIDCR/NIDCD Genomics and Computational Biology Core (ZIC DC000086), and the NIDCR Combined Technical Research Core (ZIC DE000729-09). Cary S Agler and Kimon Divaris are supported by NIDCR U01DE025046. Matthew J Flick is supported by NIH R01DK112778 and NIH R01CA211098. Francis J Castellino is supported by NIH R01HL013423-43A1.

## References

1. Belkaid, Y. and O.J. Harrison, Homeostatic Immunity and the Microbiota. Immunity, 2017. 46(4): p. 562–576.

2. Moutsopoulos, N.M. and J.E. Konkel, Tissue-Specific Immunity at the Oral Mucosal Barrier. Trends Immunol, 2018. 39(4): p. 276–287.

3. Tefs, K., et al., Molecular and clinical spectrum of type I plasminogen deficiency: A series of 50 patients. Blood, 2006. 108(9): p. 3021–6.

4. Schuster, V., B. Hugle, and K. Tefs, Plasminogen deficiency. J Thromb Haemost, 2007. 5(12): p. 2315–22.

5. Frimodt-Moller, J., Conjunctivitis ligneosa combined with a dental affection. Report of a case. Acta Ophthalmol (Copenh), 1973. 51(1): p. 34–8.

6. Kurtulus Waschulewski, I., et al., Immunohistochemical analysis of the gingiva with periodontitis of type I plasminogen deficiency compared to gingiva with gingivitis and periodontitis and healthy gingiva. Arch Oral Biol, 2016. 72: p. 75–86.

7. Ertas, U., N. Saruhan, and O. Gunhan, Ligneous periodontitis in a child with plasminogen deficiency. Niger J Clin Pract, 2017. 20(12): p. 1656–1658.

8. Neering, S.H., et al., Periodontitis associated with plasminogen deficiency: a case report. BMC Oral Health, 2015. 15: p. 59.

9. El-Darouti, M., et al., Ligneous conjunctivitis with oral mucous membrane involvement and decreased plasminogen level. Pediatr Dermatol, 2009. 26(4): p. 448–51.

10. Darveau, R.P., Periodontitis: a polymicrobial disruption of host homeostasis. Nat Rev Microbiol, 2010. 8(7): p. 481–90.

11. Hajishengallis, G., Periodontitis: from microbial immune subversion to systemic inflammation. Nat Rev Immunol, 2015. 15(1): p. 30–44.

12. Bugge, T.H., et al., Plasminogen deficiency causes severe thrombosis but is compatible with development and reproduction. Genes Dev, 1995. 9(7): p. 794–807.

13. Bugge, T.H., et al., Loss of fibrinogen rescues mice from the pleiotropic effects of plasminogen deficiency. Cell, 1996. 87(4): p. 709–19.

14. Sulniute, R., et al., Plasmin is essential in preventing periodontitis in mice. Am J Pathol, 2011. 179(2): p. 819–28.

15. Drew, A.F., et al., Ligneous conjunctivitis in plasminogen-deficient mice. Blood, 1998. 91(5): p. 1616–24.

16. Luyendyk, J.P., J.G. Schoenecker, and M.J. Flick, The multifaceted role of fibrinogen in tissue injury and inflammation. Blood, 2019. 133(6): p. 511–520.

17. Flick, M.J., X. Du, and J.L. Degen, Fibrin(ogen)-alpha M beta 2 interactions regulate leukocyte function and innate immunity in vivo. Exp Biol Med (Maywood), 2004. 229(11): p. 1105–10.

18. Flick, M.J., et al., Leukocyte engagement of fibrin(ogen) via the integrin receptor alphaMbeta2/Mac-1 is critical for host inflammatory response in vivo. J Clin Invest, 2004. 113(11): p. 1596–606.

19. Eskan, M.A., et al., The leukocyte integrin antagonist Del-1 inhibits IL-17-mediated inflammatory bone loss. Nat Immunol, 2012. 13(5): p. 465–73.

20. Liang, S., et al., Periodontal inflammation and bone loss in aged mice. J Periodontal Res, 2010. 45(4): p. 574–8.

21. Hajishengallis, E. and G. Hajishengallis, Neutrophil homeostasis and periodontal health in children and adults. J Dent Res, 2014. 93(3): p. 231–7.

22. Hajishengallis, G., et al., Neutrophil homeostasis and inflammation: novel paradigms from studying periodontitis. J Leukoc Biol, 2015. 98(4): p. 539–48.

23. Munz, M., et al., Meta-analysis of genome-wide association studies of aggressive and chronic periodontitis identifies two novel risk loci. Eur J Hum Genet, 2019. 27(1): p. 102–113.

24. Schaefer, A.S., et al., Genetic evidence for PLASMINOGEN as a shared genetic risk factor of coronary artery disease and periodontitis. Circ Cardiovasc Genet, 2015. 8(1): p. 159–67.

25. Medved, L., G. Tsurupa, and S. Yakovlev, Conformational changes upon conversion of fibrinogen into fibrin. The mechanisms of exposure of cryptic sites. Ann N Y Acad Sci, 2001. 936: p. 185–204.

26. Lishko, V.K., et al., Regulated unmasking of the cryptic binding site for integrin alpha M beta 2 in the gamma C-domain of fibrinogen. Biochemistry, 2002. 41(43): p. 12942–51.

27. Flick, M.J., et al., Fibrin(ogen) exacerbates inflammatory joint disease through a mechanism linked to the integrin alphaMbeta2 binding motif. J Clin Invest, 2007. 117(11): p. 3224–35.

28. Adams, R.A., et al., The fibrin-derived gamma377-395 peptide inhibits microglia activation and suppresses relapsing paralysis in central nervous system autoimmune disease. J Exp Med, 2007. 204(3): p. 571–82.

29. Davalos, D., et al., Fibrinogen-induced perivascular microglial clustering is required for the development of axonal damage in neuroinflammation. Nat Commun, 2012. 3: p. 1227.

30. Papayannopoulos, V., et al., Neutrophil elastase and myeloperoxidase regulate the formation of neutrophil extracellular traps. J Cell Biol, 2010. 191(3): p. 677–91.

31. Brinkmann, V., et al., Neutrophil extracellular traps kill bacteria. Science, 2004. 303(5663): p. 1532–5.

32. Branzk, N., et al., Neutrophils sense microbe size and selectively release neutrophil extracellular traps in response to large pathogens. Nat Immunol, 2014. 15(11): p. 1017–25.

33. Amulic, B., et al., Neutrophil function: from mechanisms to disease. Annu Rev Immunol, 2012. 30: p. 459–89.

34. Kolaczkowska, E. and P. Kubes, Neutrophil recruitment and function in health and inflammation. Nat Rev Immunol, 2013. 13(3): p. 159–75.

35. Ley, K., et al., Neutrophils: New insights and open questions. Sci Immunol, 2018. 3(30).

36. Sollberger, G., D.O. Tilley, and A. Zychlinsky, Neutrophil Extracellular Traps: The Biology of Chromatin Externalization. Dev Cell, 2018. 44(5): p. 542–553.

37. Castanheira, F.V.S. and P. Kubes, Neutrophils and NETs in modulating acute and chronic inflammation. Blood, 2019. 133(20): p. 2178–2185.

38. Chang, M.C., et al., Butyrate induces reactive oxygen species production and affects cell cycle progression in human gingival fibroblasts. J Periodontal Res, 2013. 48(1): p. 66–73.

39. Bax, B.E., et al., Stimulation of osteoclastic bone resorption by hydrogen peroxide. Biochem Biophys Res Commun, 1992. 183(3): p. 1153–8.

40. Ha, H., et al., Reactive oxygen species mediate RANK signaling in osteoclasts. Exp Cell Res, 2004. 301(2): p. 119–27.

41. Khandpur, R., et al., NETs are a source of citrullinated autoantigens and stimulate inflammatory responses in rheumatoid arthritis. Sci Transl Med, 2013. 5(178): p. 178ra40.

42. Carmona-Rivera, C., et al., Synovial fibroblast-neutrophil interactions promote pathogenic adaptive immunity in rheumatoid arthritis. Sci Immunol, 2017. 2(10).

43. Kenny, E.F., et al., Diverse stimuli engage different neutrophil extracellular trap pathways. Elife, 2017. 6.

44. Husemann, J., et al., CD11b/CD18 mediates production of reactive oxygen species by mouse and human macrophages adherent to matrixes containing oxidized LDL. Arteriosclerosis Thrombosis and Vascular Biology, 2001. 21(8): p. 1301–1305.

45. Johnson, C.M., et al., Integrin Cross-Talk Regulates the Human Neutrophil Response to Fungal beta-Glucan in the Context of the Extracellular Matrix: A Prominent Role for VLA3 in the Antifungal Response. J Immunol, 2017. 198(1): p. 318–334.

46. Eke, P.I., et al., Prevalence of periodontitis in adults in the United States: 2009 and 2010. J Dent Res, 2012. 91(10): p. 914–20.

47. Silva, L.M., L. Brenchley, and N.M. Moutsopoulos, Primary immunodeficiencies reveal the essential role of tissue neutrophils in periodontitis. Immunol Rev, 2019. 287(1): p. 226–235.

48. Moutsopoulos, N.M., et al., Interleukin-12 and Interleukin-23 Blockade in Leukocyte Adhesion Deficiency Type 1. N Engl J Med, 2017. 376(12): p. 1141–1146.

49. Hajishengallis, G., et al., Immune and regulatory functions of neutrophils in inflammatory bone loss. Semin Immunol, 2016. 28(2): p. 146–58.

50. Matthews, J.B., et al., Hyperactivity and reactivity of peripheral blood neutrophils in chronic periodontitis. Clin Exp Immunol, 2007. 147(2): p. 255–64.

51. Matthews, J.B., et al., Neutrophil hyper-responsiveness in periodontitis. J Dent Res, 2007. 86(8): p. 718–22.

52. Nicu, E.A., et al., Characterization of oral polymorphonuclear neutrophils in periodontitis patients: a case-control study. BMC Oral Health, 2018. 18(1): p. 149.

53. White, P.C., et al., Neutrophil Extracellular Traps in Periodontitis: A Web of Intrigue. J Dent Res, 2016. 95(1): p. 26–34.

54. Dutzan, N., et al., A dysbiotic microbiome triggers TH17 cells to mediate oral mucosal immunopathology in mice and humans. Sci Transl Med, 2018. 10(463).

55. Ryu, J.K., et al., Fibrin-targeting immunotherapy protects against neuroinflammation and neurodegeneration. Nat Immunol, 2018. 19(11): p. 1212–1223.

56. Schacher, B., et al., Aggregatibacter actinomycetemcomitans as indicator for aggressive periodontitis by two analysing strategies. J Clin Periodontol, 2007. 34(7): p. 566–73.

57. Overton, N.L., et al., Genetic susceptibility to severe asthma with fungal sensitization. Int J Immunogenet, 2017. 44(3): p. 93–106.

58. Suh, T.T., et al., Resolution of spontaneous bleeding events but failure of pregnancy in fibrinogen-deficient mice. Genes Dev, 1995. 9(16): p. 2020–33.

59. Carmeliet, P., et al., Physiological consequences of loss of plasminogen activator gene function in mice. Nature, 1994. 368(6470): p. 419–24.

60. Coxon, A., et al., A novel role for the beta 2 integrin CD11b/CD18 in neutrophil apoptosis: a homeostatic mechanism in inflammation. Immunity, 1996. 5(6): p. 653–66.

61. Iwaki, T., et al., The generation and characterization of mice expressing a plasmin-inactivating active site mutation. J Thromb Haemost, 2010. 8(10): p. 2341–4.

62. Bugge, T.H., et al., Urokinase-type plasminogen activator is effective in fibrin clearance in the absence of its receptor or tissue-type plasminogen activator. Proc Natl Acad Sci U S A, 1996. 93(12): p. 5899–904.

63. Wilensky, A., et al., Three-Dimensional Quantification of Alveolar Bone Loss in Porphyromonas gingivalis-Infected Mice Using Micro-Computed Tomography. Journal of Periodontology, 2005. 76(8): p. 1282–1286.

64. Lendrum, A.C., et al., Studies on the character and staining of fibrin. J Clin Pathol, 1962. 15: p. 401–13.

65. Dutzan, N., et al., Isolation, Characterization and Functional Examination of the Gingival Immune Cell Network. J Vis Exp, 2016(108): p. 53736.

66. Prasad, J.M., et al., Mice expressing a mutant form of fibrinogen that cannot support fibrin formation exhibit compromised antimicrobial host defense. Blood, 2015. 126(17): p. 2047–58.

67. Swamydas, M. and M.S. Lionakis, Isolation, purification and labeling of mouse bone marrow neutrophils for functional studies and adoptive transfer experiments. J Vis Exp, 2013(77): p. e50586.

68. Divaris, K., et al., Exploring the genetic basis of chronic periodontitis: a genome-wide association study. Hum Mol Genet, 2013. 22(11): p. 2312–24.

69. Beck, J.D., et al., In search of appropriate measures of periodontal status: The Periodontal Profile Phenotype (P(3)) system. J Periodontol, 2018. 89(2): p. 166–175.

70. Morelli, T., et al., Derivation and Validation of the Periodontal and Tooth Profile Classification System for Patient Stratification. J Periodontol, 2017. 88(2): p. 153–165.

71. Darveau, R.P., G. Hajishengallis, and M.A. Curtis, Porphyromonas gingivalis as a potential community activist for disease. J Dent Res, 2012. 91(9): p. 816–20.

72. Segre, A.V., et al., Common inherited variation in mitochondrial genes is not enriched for associations with type 2 diabetes or related glycemic traits. PLoS Genet, 2010. 6(8).

73. Pruim, R.J., et al., LocusZoom: regional visualization of genome-wide association scan results. Bioinformatics, 2010. 26(18): p. 2336–7.

