## Supplementary Figures for "Fibrin is a critical regulator of neutrophil effector function at mucosal barrier sites"

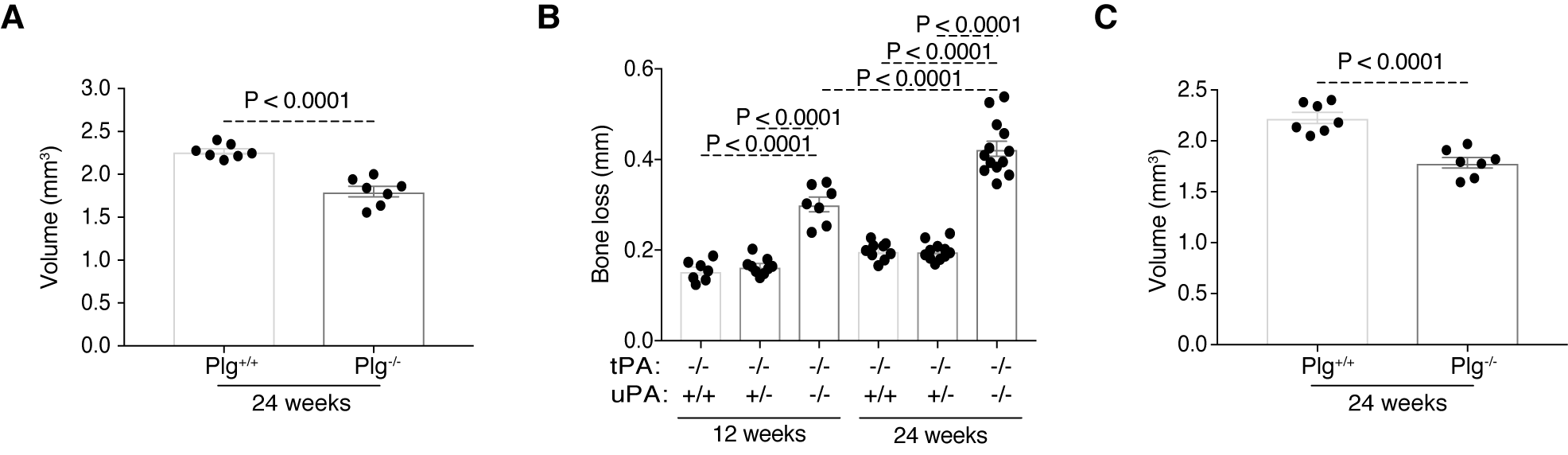

**Supplementary Figure 1 (related to Figure 1): Activation of plasminogen is essential for maintaining periodontal health.** (A) μCT analysis of alveolar bone volume (mm^3^) in *Plg^-/-^* mice compared to the wild-type controls. (B) bone loss measurement in *Plat^-/-^;Plau^+/+^, Plat^-/-^;Plau^+/-^* and *Plat^-/-^;Plau^-/-^* mice maxillae.

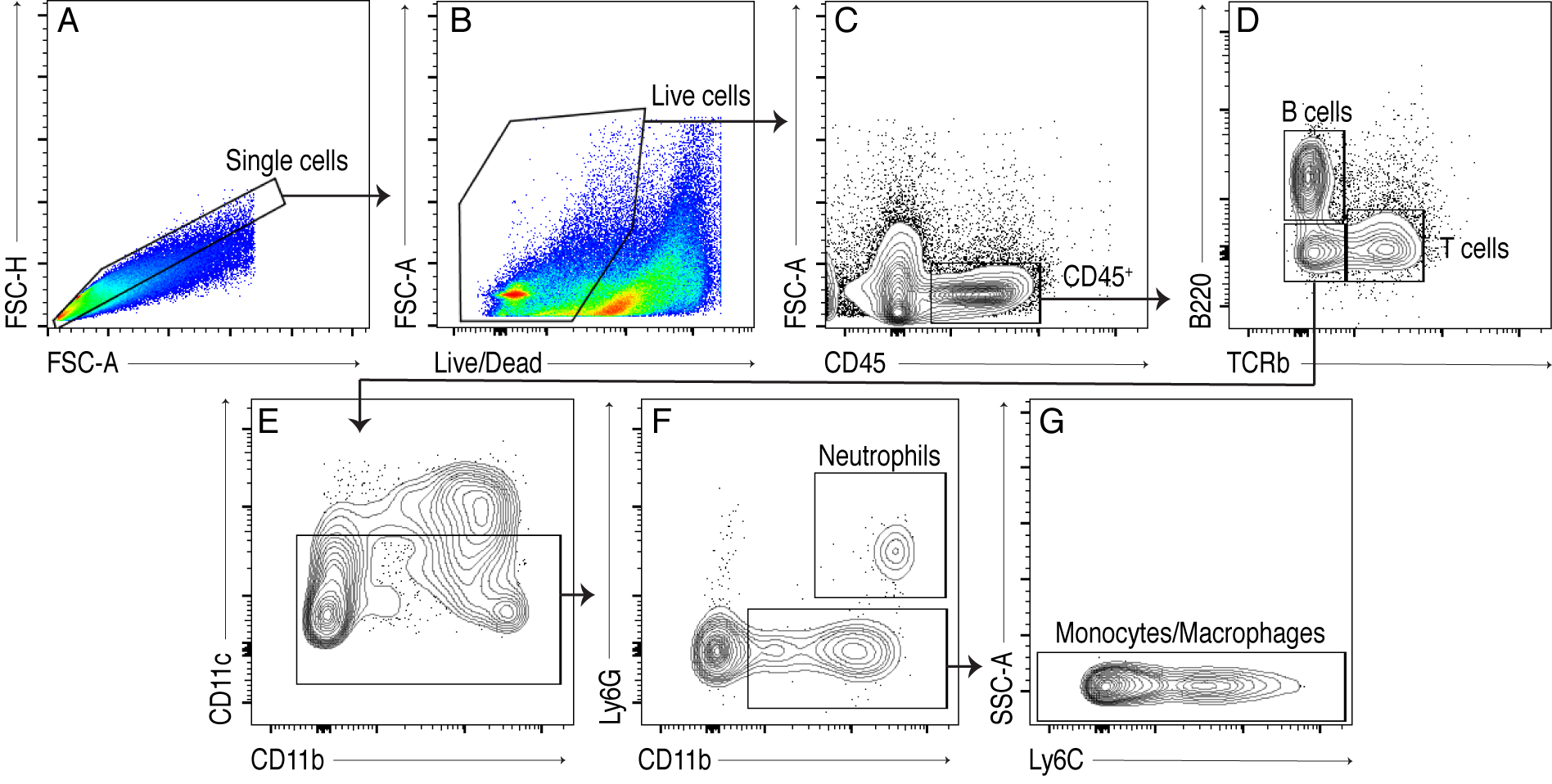

**Supplementary Figure 2 (related to Figure 2): Gating strategy for flow cytometry analysis of immune cell populations.** (A) Single cells were determined by gating for FSC-A and FSC-H. (B) Live cells were gated for (C) CD45+ cells. (D) T cell and B cell populations were determined by the presence of TCRb and B220 markers, respectively. The TCRb- B220- population was further gated to select (E) CD11c Low-Med cells. (F) Neutrophils were characterized as CD11b+ Ly6G+ cells. CD11b+ Ly6G- cells were further gated for (G) Ly6C and SSC to select the SSC-Low monocyte/macrophage population.

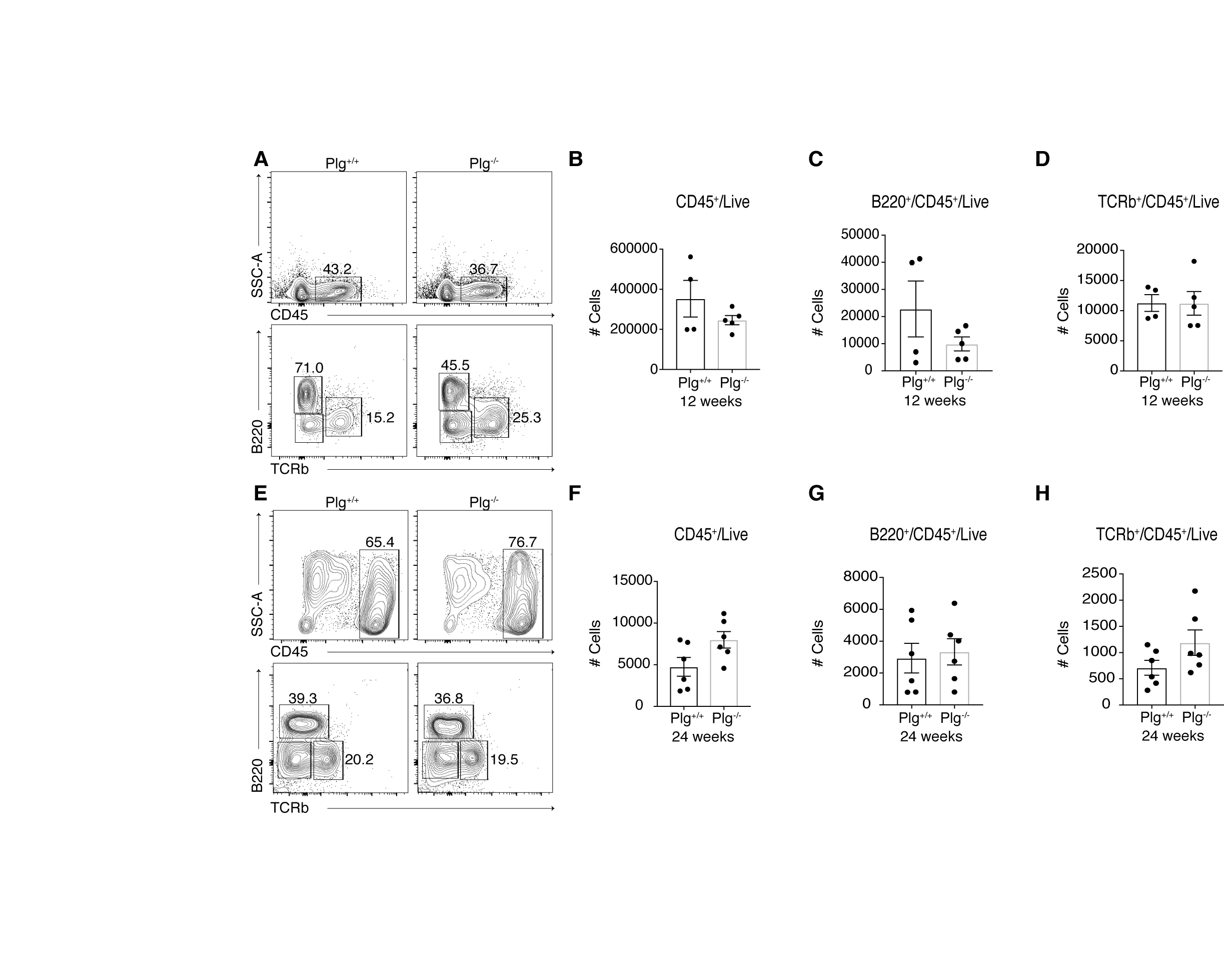

**Supplementary Figure 3 (related to Figure 2 and 3): T cells and B cells are not significantly changed in Plg-deficient mice gingivae.** (A) Flow cytometry analysis of 12-weeks old mouse gingival tissues. Contour plots show changes in total hematopoietic cells (Live/CD45+) and T (Live/CD45+TCRb+) and B (Live/CD45+/B220+) cell populations in percentages. Counts of (B) total hematopoietic cells; (C) B cells; and (D) T cells 12-week-old in *Plg^+/+^* and *Plg^-/-^* mouse gingivae. (E) Flow cytometry analysis of 24-week-old mouse gingival tissues. Contour plots show changes in total hematopoietic cells and T and B cell populations in percentages. Counts of (F) total hematopoietic cells; (G) B cells; and (H) T cells in 24-week-old *Plg^+/+^* and *Plg^-/-^* mouse gingivae.

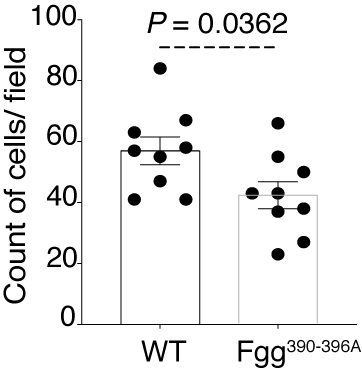

**Supplementary Figure 4 (Related to Figure 4):** **Fibrin(ogen)-neutrophil interaction through α_M_β_2_-binding.** Mouse neutrophil binding to *Fgg^390-396A/390-396A^* fibrin compared to wild-type fibrin.

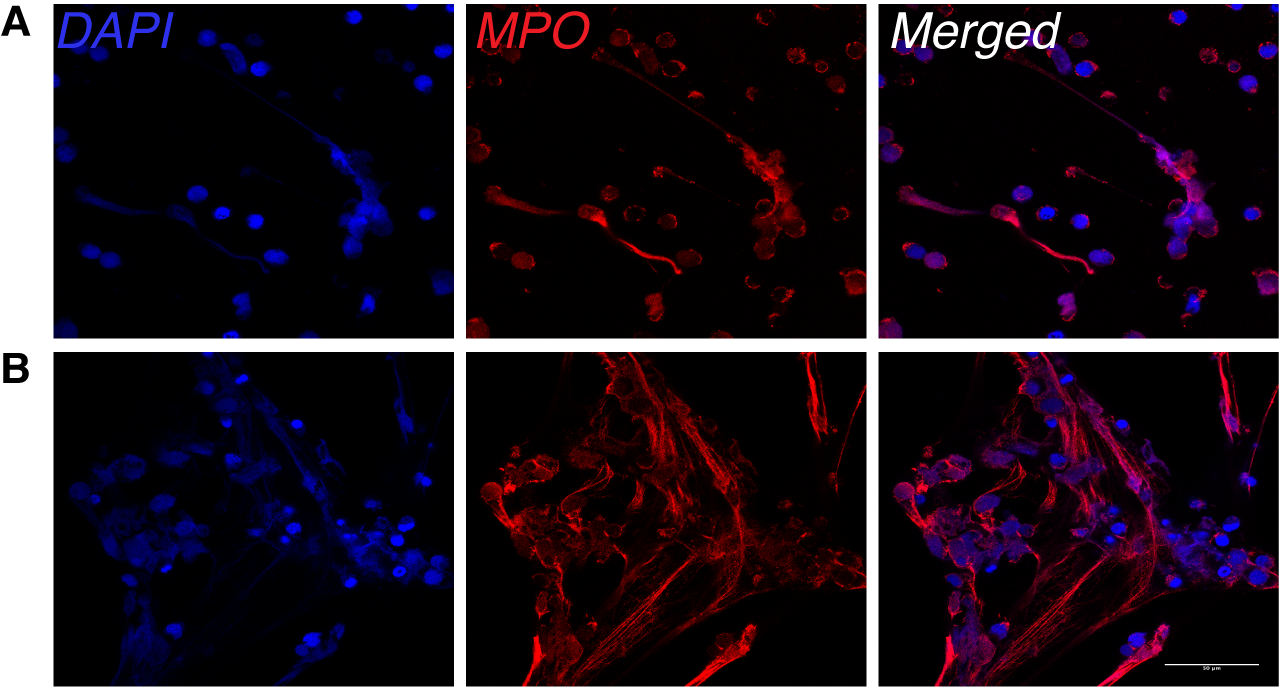

**Supplementary Figure 5 (Related to Figure 4): Visualization of NETing neutrophils.** Human neutrophils plated on wild-type fibrin stained for DAPI (blue) and MPO (red) showing (A) intracellular MPO staining in intact neutrophils and (B) externalized MPO and DAPI in NETing neutrophils (Scale bar = 50 μm).

**Supplementary video (Related to Figure 4):** **NETosis with wild-type or Fgg^390-396A^ fibrin.** Random video for a time lapse series showing live neutrophils (Hoechst:blue), and those undergoing either NETosis (Red overlay) or apoptosis (Cytox Green:green) after stimulation with 15 nM PMA plated on either wild-type or Fgg^390-396A^ fibrin.

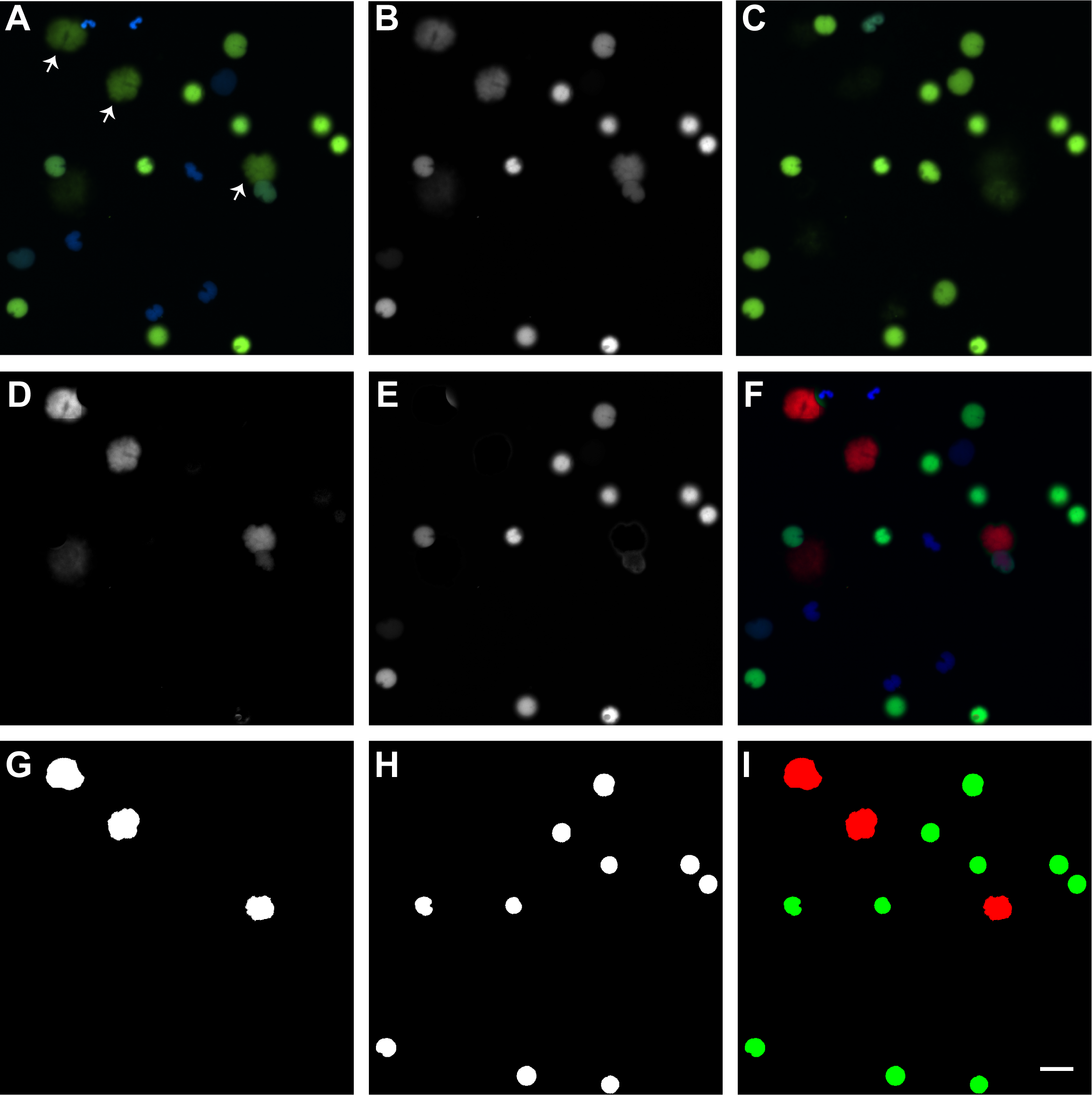

**Supplementary Figure 6 (Related to Figure 4): NETosis and apoptosis image analysis scheme.** (A) Random image for a time lapse series showing live neutrophils (Hoechst:blue), and those undergoing either NETosis or apoptosis (Cytox Green:green) after stimulation with 15 nM PMA. Arrows indicate neutrophils undergoing NETosis. (B) Cytox Green channel from A. (C) Last image of a representative 5-hour time lapse series showing only apoptotic cells (green). Note the lack of cells undergoing NETosis. This image is subtracted from the panel B to yield the image shown in panel (D) that shows the “NETosis-only fraction”. (E) Subsequent subtraction of the “NETosis-only fraction” produces an “Apoptosis-only fraction”. (F) Color overlay shows live neutrophils (blue), apoptotic neutrophils (green) and neutrophil that underwent NETsosis (red). (G) Otsu threshold mask applied to “NETosis-only fraction” in panel D. (H) Otsu threshold mask applied to “Apoptosis-only fraction” in panel E. (I) Overlay of NETosis (red) and apoptosis (green) masks. Scale bars: 25 μm.

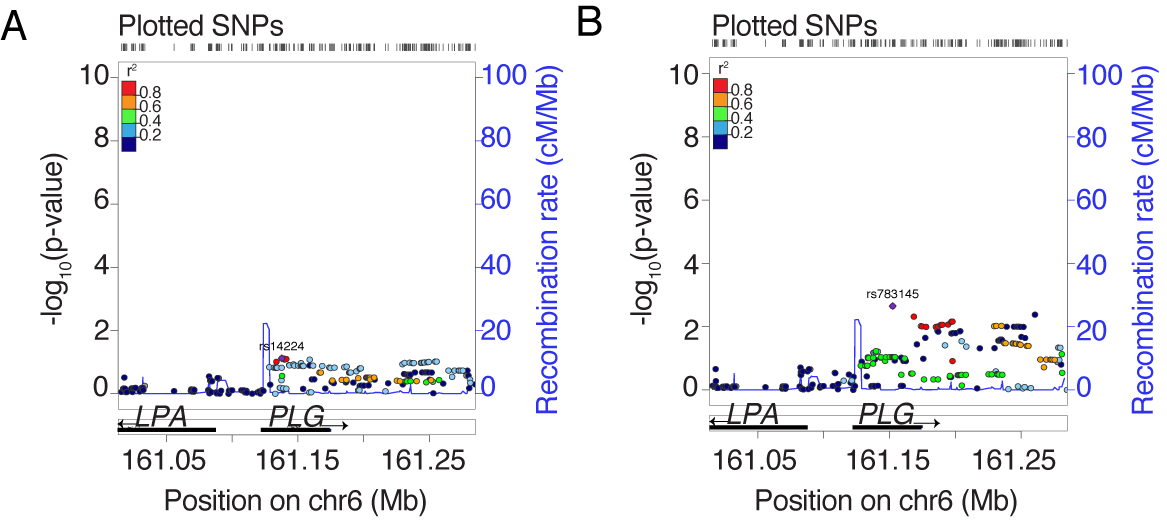

**Supplementary Figure 7 (related to Figure 6): Regional association plots of *PLG*** with mild disease (A) and severe gingival inflammation (B) in the Dental Atherosclerosis Risk In Communities study (D-ARIC) population.
